# The population genetics of adaptation through copy-number variation in a fungal plant pathogen

**DOI:** 10.1101/2021.12.10.472168

**Authors:** Luzia Stalder, Ursula Oggenfuss, Norfarhan Mohd-Assaad, Daniel Croll

**Affiliations:** Laboratory of Evolutionary Genetics, Institute of Biology, University of Neuchâtel, 2000 Neuchâtel, Switzerland; Plant Pathology, Institute of Integrative Biology, ETH, Zurich, 8092 Zurich, Switzerland; School of Biosciences and Biotechnology, Faculty of Science and Technology, Universiti Kebangsaan Malaysia, 43600 Bangi, Selangor, Malaysia

**Keywords:** copy number variation, adaptation, plant pathogen, fungi

## Abstract

Microbial pathogens can rapidly adapt to changing environments such as the application of pesticides or host resistance. Copy number variations (CNV) are a major source of adaptive genetic variation for recent adaptation. Here, we analyze how a major fungal pathogen of barley, *Rhynchosporium commune*, has adapted to host environment, fungicide and temperature challenges. We screen the genomes of 126 isolates sampled across a worldwide set of populations and identify a total of 7’879 gene duplications and 116 gene deletions. Most gene duplications result from segmental chromosomal duplications. We find that genes showing recent gains or losses are enriched in functions related to host exploitation (*i.e.* effectors and cell wall degrading enzymes). We perform a phylogeny-informed genome-wide association study (GWAS) and identify 191 copy-number variants associated with different pathogenesis and temperature related traits, including a large segmental duplication of *CYP51A* that has contributed to the emergence of azole resistance. Additionally, we use a genome-wide SNP dataset to replicate the GWAS and contrast it with the CNV-focused analysis. We find that frequencies of adaptive CNV alleles show high variation among populations for traits under strong selection such as fungicide resistance. In contrast, adaptive CNV alleles underpinning temperature adaptation tend to be near fixation. Finally, we show that transposable elements are important drivers of recent gene copy-number variation. Loci showing signatures of recent positive selection are enriched in miniature inverted repeat transposons. Our findings show how extensive segmental duplications create the raw material for recent adaptation in global populations of a fungal pathogen.

## INTRODUCTION

The incidence of emerging pathogens has been increasing and presents a major threat to worldwide food security today (Fisher et al. 2012, 2018; Fones et al. 2020; Brown et al. 2012). Among pests, fungi cause the most crop devastation, accounting for ∼30% of perennial yield losses worldwide (Fisher et al. 2018). Major challenges come from the ability of fungal pathogens to rapidly adapt to environmental changes such as the application of fungicides (Fisher et al. 2018). Point mutations and structural variants, such as copy number variants (CNV), inversions or translocations provide genetic raw material for selection to act upon. In particular, gene duplications or deletions are a major source of phenotypic variation driving species adaptation (Wellenreuther et al. 2019; Mérot et al. 2020). In the case of pathogens, the loss of genes that are recognized by the immune system of the host can be highly beneficial (Olson 1999; Morris, Lenski, and Zinser 2012; Hartmann and Croll 2017). In contrast, duplications of genes targeted by fungicides have been shown to confer resistance in several species (Coste et al. 2007; Sionov et al. 2010; Zhang et al. 2019; Morschhäuser 2016; Steenwyk and Rokas 2018; Sánchez-Torres 2021; Leroux and Walker 2013; Ma et al. 2006). Yet, despite their high adaptive potential, systematic population-wide analyses of adaptive CNV have been scarce (Mérot et al. 2020).

The high adaptive potential of CNV comes from their high genomic abundance, as well as from their manyfold phenotypic effects (Tang and Amon 2013). In adaptive evolution experiments with nutrient-deprivation schemes, genes that facilitate the uptake and/or metabolism of the limiting factor are upregulated often through gene duplication (Tang and Amon 2013). For example, glucose limitation in *Saccharomyces cerevisiae* cultures has resulted in gene duplications encoding glucose transporters, while sulfate limitation resulted in a gene duplication encoding a high-affinity sulfate transporter (Mishra and Whetstine 2016). In line with these observations, CNV of genes targeted by fungicides have been repeatedly associated with resistance (Coste et al. 2007; Sionov et al. 2010; Zhang et al. 2019; Morschhäuser 2016; Steenwyk and Rokas 2018; Sánchez-Torres 2021; Leroux and Walker 2013; Ma et al. 2006). Azoles are the most used fungicides both in agricultural practices and clinical settings to treat fungal infections (Cools and Fraaije 2013; Azevedo et al. 2015). The dominant mechanisms to gain azole resistance are non-synonymous mutations as well as CNV of the gene *CYP51* (*ERG11* in yeast), a member of the cytochrome P450 family and the direct molecular target of azoles. Human pathogenic fungi including members of the *Candida* and *Cryptococcus* geni have repeatedly acquired azole resistance (Coste et al. 2007; Todd and Selmecki 2020; Sionov et al. 2010; Morschhäuser 2016). During infection, *Candida* species show loss-of-heterozygosity allowing for more efficient selection as quasi-haploids (Todd and Selmecki 2020). Similarly, also many crop pathogens have rapidly gained azole resistance through mutations or duplications of *CYP51* (Lucas, Hawkins, and Fraaije 2015). A particularly interesting case is the barley pathogen *Rhynchosporium commune*. In addition to adaptive mutations in *CYP51*, the *R. commune* genome encodes a total of three *CYP51* paralogues including *CYP51A*, *CYP51B* and *CYP51A-p* (a loss of function copy of *CYP51A*) (Brunner et al. 2016; Hawkins et al. 2014). The *CYP51A* presence/absence polymorphism in *R. commune* is strongly correlated with azole resistance on a global scale (Brunner et al. 2016). Populations from the United Kingdom had very low *CYP51A* frequencies between 1890 and 1985 and then the *CYP51A* frequency increased very rapidly, presumably as a response to increased azole treatments (Hawkins et al. 2014). CNVs in *R. commune* have also impacted genes contributing to virulence, including the genes encoding for the necrosis-inducing proteins *NIP1*, *NIP2* and *NIP3* (Penselin et al. 2016). *NIP1* is recognized by the barley resistance gene *Rrs1*, impeding infection in a gene-for-gene model (Schürch et al. 2004; Kirsten et al. 2012; Rohe et al. 1995). Consequently, the *NIP1* gene has been undergoing several gene duplication events with some predating speciation (Mohd□ Assaad, McDonald, and Croll 2019). In addition, the adaptive deletion of the *NIP1* gene allows *R. commune* to infect barley cultivars carrying the cognate resistance factor *Rrs1*(Mohd□ Assaad, McDonald, and Croll 2019).

In eukaryotic microbial pathogens, adaptive CNVs are generated both during mitosis and meiosis (Hastings et al. 2009). During meiosis, nonhomologous recombination between specific chromosomal regions can cause the duplication and deletion of sequences among progeny (Hastings et al. 2009). In absence of sexual reproduction, non-allelic homologous recombination during mitosis or aneuploidy are critical to generate genetic variability. Importantly, generation of CNV is often driven by transposable elements (TEs), ubiquitous repetitive DNA sequences that can move from one location in the genome to another (Wells and Feschotte 2020; Bourque et al. 2018; Feschotte 2008). TEs can either passively facilitate structural variation by increasing the likelihood of non-homologous recombination, or actively insert into new regions of the genome and create CNVs in populations (Wells and Feschotte 2020; Bourque et al. 2018; Feschotte 2008). The recent advent of genome analyses has allowed to establish mechanisms giving rise to structural variation (Mérot et al. 2020). However, how specific genomic features in microbial pathogens influence the creation and population structure influences the trajectory of adaptive CNVs remains largely unknown (Mérot et al. 2020).

Here we investigate how the barley pathogen *R. commune* has adapted to fungicide applications, host resistance and different temperature regimes at a global scale. Using phylogeny-informed genome- wide association mapping, we identify a broad set of CNV loci likely contributing to variation in life- history traits, including large segmental duplications of the fungicide resistance locus encoding *CYP51A*. We contrast the findings on CNV associations with mapping outcomes based on genome- wide single nucleotide polymorphisms (SNPs). We find that CNVs are enriched for gene functions related to host colonization and environmental adaptation. We show that CNVs associated with phenotypic trait variation are close to specific TE categories. More broadly, CNVs under recent positive selection are over-represented for loci with TE insertions of specific elements. Together, our findings highlight the population genetic context and genomic environment of CNV-based adaptation in populations.

## MATERIAL AND METHODS

### FIELD COLLECTION OF *R. COMMUNE*

We analyzed 126 isolates of *R. commune* collected from naturally infected barley fields. Populations from nine different countries and four different continents were obtained: New Zealand (NZ), Australia (AU), Ethiopia (ET), Switzerland (CH), Spain (SP), Norway (NO), Finland (FI), Iceland (IS) and USA (US). All isolates were previously characterized using microsatellite markers for population genetics studies, and whole-genome sequencing data is available for 12-14 isolates per population (Mohd□ Assaad, McDonald, and Croll 2016; Tryggvi S. Stefansson, Mcdonald, and Willi 2013; Tryggvi S. Stefansson, McDonald, and Willi 2014).

### FUNGICIDE SENSITIVITY ASSAY

Fungicide sensitivity was assessed as previously described (Mohd□ Assaad, McDonald, and Croll 2016). Isolates were taken from -80 °C storage by plating on lima bean agar plates (LBA; 60 g/L lima bean, 12 g/L Bacteriological Agar, 50 mg/L kanamycin). After 10 days at 18 °C in darkness, spores were harvested by flooding the plate with sterile water, aided by scraping with a microscope slide and filtering through two layers of sterile cheesecloth. The quantification of spore densities was done using KOVATM GlassticTM Slides that were adjusted to a concentration of 105 spores m/L. Cyproconazole (Syngenta Inc., Stein, Switzerland) was serially diluted to achieve final concentrations of 375, 37.5, 3.75, 0.375, 0.0375, 0.00375 and 0.000375 mg/L. 150 µl of Synthetic Nutrient Broth (SNB; 0.2% (w/v) KH2PO4, 0.2% (w/v) KNO3, 0.1% (w/v) MgSO4.7H2O, 0.1% (w/v) KCl and 0.2% (w/v) sucrose) together with cyproconazole was prepared in each well and 50 µl of spore suspension or sterile water as a negative control was added. Each isolate was tested in four technical replicates. Parafilm-covered plates were incubated in darkness at 18 °C with 80% humidity for 7 days. Then, 20 µ l of filter-sterilized 550 µM resazurin (RZ; Sigma-Aldrich) dissolved in phosphate-buffered saline solution (PBS, pH 7.4) was added to each well, followed by 24 h incubation. RZ measures the metabolic activity of fungal growth, hence the reduction in RZ was used as a measure of fungal hyphal growth in the microtiter plate assay (Vega et al. 2012). The reduction in RZ was measured using a SpectraMax_i3 Multi-Mode Microplate Reader Platform with the SpectraMax- MiniMaxTM Imaging Cytometer (Molecular Devices, Sunnyvale, CA) at excitation and emission wavelengths of 565 nm and 590 nm, respectively. The dose-response curve was fitted for each assay individually using R with the four-parameter log-logistic model implemented in the DRC package (Ritz et al. 2015).

### VIRULENCE ASSAY

Virulence was assessed as previously described (T. S. Stefansson et al. 2014). Spore solutions were prepared from -80°C storage as described above. The quantification of spore density was done using a haemocytometer and concentrations were adjusted to 10^6^ spores ml^-1^. Spore solution volume was adjusted to 5 ml and 5 µ l Tween 20 was added. Virulence was measured on the spring barley cultivar Beatrix (Viskosa 9 Pasadena, Saaten Union, breeders’ Reference NS01/2449). Pots with soil and four seeds each were placed in a single greenhouse chamber, where plants were grown at 18°C during the day and 15°C during the night. Day length was set to 14 h and humidity was set at 60%. Watering occurred from the bottom. After emergence of the second leaf (∼12 days), each plant was inoculated until run-off in a sterile chamber. A spore suspension of 5 ml was used per isolate and each isolate was replicated three times. Plants were dried for 15-20 min and then arranged randomly in mobile humidity chambers that were placed in the growth chamber. Relative humidity was kept at 100% for 48 h with the temperature and day length conditions as described above. The plants were then randomly arranged in a single greenhouse chamber and the relative humidity was set to 60%. Scoring for symptoms was done for 1 week beginning 9 days after inoculation. Fifteen days after inoculation, the second leaf on all plants was cut at the leaf base and mounted onto a light-blue paper sheet (size A4). Using a Canon EOS 60D camera and a 50 mm lens, digital images were taking in a darkroom. On each side of the mounted leaves, a fixed light source (Philips PF 319 E/44, 150 W) was placed with a distance of 30 cm. The percentage of leaf area covered by lesions was measured for each leaf using the image analysis software APS ASSESS (Lamari 2002). The total number of isolates included in the analysis was 114 (12 isolates missing). Analysis was conducted using the manual panel and thresholds were set by hue in APS ASSESS. Virulence was logit-transformed prior to analysis due to a zero inflation.

### GROWTH RATES AND COLONY MELANIZATION AT DIFFERENT TEMPERATURES

Growth rates at 12 °C, 18 °C and 22 °C were assessed as previously described (Tryggvi S. Stefansson, Mcdonald, and Willi 2013). Temperature sensitivity was calculated as the standard deviation of the growth rate at 12 °C, 18 °C and 22 °C. Hence, a lower standard deviation indicates higher growth resilience to temperature variation. Melanization was measured at 12 °C, 18 °C and 22 °C in the same setting as described above. The digital images of fungal colonies were analyzed for melanization using automated image analysis based on ImageJ (Schneider, Rasband, and Eliceiri 2012). The method was identical to the one described for *Zymoseptoria tritici*, with the only difference that the gray scale (GS) used to measure melanization was inverted using the formula 255-GS so that 255 equals black and 0 equals white (Lendenmann, Croll, and McDonald 2015). Hence, higher values indicate higher levels of melanization.

### DNA EXTRACTION, GENOME SEQUENCING, READ MAPPING

DNA extraction and sequencing was done as previously described (Mohd□ Assaad, McDonald, and Croll 2016). Genomic DNA was extracted from mycelium cultured on Potato Dextrose Broth (PDB) using DNeasy Plant Mini Kits (Qiagen) according to the manufacturer’s instructions. Paired-end reads of 125 bp each were sequenced on a Illumina HiSeq 2000 platform with an insert size of approximately 500 bp. The sequencing was performed at the Quantitative Genomics Facility of the D-BSSE at the ETH Zurich, Switzerland. The sequencing data are available at the NCBI Sequence Read Archive under the BioProject accession number PRJNA327656. Reads were screened for adapter contamination and trimmed for sequencing quality using Trimmomatic v. 0.32 with the following settings: trailing = 10, sliding-window = 4:10 and minlen = 50 (Bolger, Lohse, and Usadel 2014). Trimmed reads were aligned to the *R. commune* reference genome UK7 v. 2 using bowtie v. 2 to generate SAM sequence alignments. (Langmead and Salzberg 2012). The reference genome is available at the European Nucleotide Archive (http://www.ebi.co.uk/ena) under accessions FJUW01000001-FJUW01000164.

### SNP AND CNV CALLING

SNP calling was done as previously described (Mohd□ Assaad, McDonald, and Croll 2016). The HaplotypeCaller and GenotypeGVCF tools of the GATK 3.3-0 suite were used (McKenna et al. 2010). SNPs with a phred-scaled quality score lower than 500 were removed and SNPs were retained if they satisfied the following conditions: -2 < BaseQualityRankSumTest < 2, -2 < ReadPosRankSumTest < 2, RMSMappingQuality < 30, -2 < MappingQualityRankSumTest < 2, QualByDepth > 20 and FisherStrand < 10. The SNP data set was filtered to retain only SNPs with a genotyping rate >90% and a minor allele frequency (MAF) >5%.

To call CNVs, the software CNVnator was used (Abyzov et al. 2011). Statistical analysis of short read coverage along the genome was performed to identify CNVs in all 126 sequenced isolates compared to the UK7 reference genome. For each isolate, CNV events were assessed in 100 bp bins as recommended. CNVs were retained if the CNV genotype *p*-value < 0.05 and q0 < 0.5. Additionally, CNVs were filtered for the normalized average read depth. Deletions with a normalized average read depth of <0.4 and duplications with normalized average read depth of >1.6 were retained.

### RNA-SEQ-ASSISTED GENE MODEL PREDICTION AND DIFFERENTIAL EXPRESSION ANALYSES

The transcription profiles of all genes were assessed using RNA-seq data generated for the reference isolate UK7. RNA-seq experiments were conducted for four conditions, two *in vitro* and two *in plantae*. *In vitro* culture conditions included growth on Luria-Bertani (LB) and Potato Dextrose Broth (PDB) media. For the *in planta* experiments, plants of the barley cultivar Beatrix (Viskosa 9 Pasadena, Saaten Union, breeders’ Reference NS01/2449) were infected, and leaves were collected at 9 and 13 days post infection (dpi). All experiments were conducted in triplicates. Total RNA was extracted using TRIzol (Invitrogen Inc.) following the manufacturer’s recommendations. RNA integrity and quantity was assessed on a Bioanalyzer 2100 (Agilent) and a Qubit fluorometer (Life Technologies). Libraries were prepared using the TruSeq stranded mRNA sample prep kit (Illumina Inc.). Total RNA was ribosome-depleted by using polyA selection and reverse-transcribed into double-stranded cDNA. Raw reads for the *in vitro* conditions are available at https://doi.org/10.5281/zenodo.5729863, and for the *in plantae* conditions at https://doi.org/10.5281/zenodo.5729968.

The software tophat v. 2.0.14 was used to align short reads to the reference genome (Trapnell, Pachter, and Salzberg 2009). To identify high-quality gene models, the reference genome was newly annotated. Intron splice site hints were generated using bam2hints, included in the AUGUSTUS v.

3.2.1 software (Stanke et al. 2006). Due to the very high RNA-sequencing depth available, intron splice hints were filtered for a minimum coverage of 20 reads to avoid an impact of spurious splice signals on gene prediction. To produce *ab initio* gene models, the BRAKER v. 1.0 pipeline combining GeneMark-ET *ab initio* gene model predictions and AUGUSTUS v. 3.2.1 (Hoff et al. 2016). GeneMark-ET was trained using the RNA-seq-based splice information as hints. AUGUSTUS was automatically trained using *ab initio* gene models that were fully supported by splice information. Finally, AUGUSTUS was used to predict gene models using both RNA-seq splice information and coding sequence hints based on exonerate protein alignments as extrinsic evidence. The gene models are available at https://doi.org/10.5281/zenodo.5730007. All newly annotated gene models for the UK7 reference genome were functionally characterized using InterProScan 5 v. 79.0 (Jones et al. 2014). All proteins predicted to be secreted using SignalP v. 4.1 were also screened for carbohydrate-active enzymes (CAZymes) using the carbohydrate-active enzyme annotation (dbCAN) release 5.0 (Yin et al. 2012; Petersen et al. 2011). Only protein hits that were predicted by Diamond, HMMER and Hotpep were retained. Orthologues to virulence and resistance associated genes were identified using BLASTP against the pathogen-host interaction database PHI-base (Urban et al. 2020). Proteins were extracted with the search terms “effector” or “resist”. Mapped reads overlapping gene models were counted using HTSeq-count v. 0.11.1, setting the matching mode to “union” and filtering out reads with an alignment quality below 10 (Anders, Pyl, and Huber 2015). Reads per kilobase of transcript per million mapped reads (RPKM) was calculated using R with the package edgeR v. 3.30.0 (Robinson, McCarthy, and Smyth 2009).

### IDENTIFICATION OF CONSERVED AND ORPHAN CNV-GENES

To identify genes affected by CNV, the percentage of overlap between CNV and genes were calculated using the bedtools v. 2.27.1 “intersect” command. (Quinlan and Hall 2010). A gene was conservatively considered as affected by a CNV if the CNV was overlapping > 80% of the gene length. The CNV overlap was defined as the sum of the overlapping deletion or duplication events. Genes affected by both deletions and duplications were excluded. To classify CNV-genes as conserved or orphan CNV-genes, we inferred orthologues of all *R. commune* genes using OrthoFinder v. 2.1.2 with default parameters (Emms and Kelly 2019). We used 68 Leotimycetes genomes including 4 *Rhynchosporium* genomes and 4 Sordariomycetes genomes for our analysis (Supplementary Fig. S4). We defined conserved *R. commune* genes as genes with orthologues in all *Rhynchosporium* species, >55 Leotimycetes and 4 Sordariomycetes. We defined *R. commune* orphan genes as genes with no orthologues in other species of the *Rhynchosporium* genus, Leotimycetes or Sordariomycetes (Supplementary Fig. S4).

### TE CONSENSUS IDENTIFICATION, CLASSIFICATION AND ANNOTATION

To obtain a consensus sequence for each TE family, RepeatModeler v. open-4.0.7 (http://www.repeatmasker.org/RepeatModeler/) was run on the *R. commune* UK7 reference genome. The classification was based on the GIRI Repbase (v. 2018) using RepeatMasker v. open-4.0.7. (Smit, Hubley, and P. 2015; Bao, Kojima, and Kohany 2015). To finalize the classification of TE consensus sequences, a manual curation with WICKERsoft was used (Breen et al. 2010). The genome was screened for copies of known consensus sequences from other fungal species with blastn filtering for sequence identity > 80% and sequence length > 80%. (Altschul et al. 1997). Flanks of 10000 bp were added and visually inspected for sequence similarity and terminal repeats with dot plots. Multiple sequence alignments were performed with 10-15 sequences using ClustalW (Thompson, Higgins, and Gibson 1994). Alignment boundaries were visually inspected and trimmed if necessary. Consensus sequences were classified according to the presence and type of terminal repeats, as well as homology of the encoded proteins based on blastx against the NCBI protein database. Consensus sequences were named according to the three-letter classification system (Wicker et al. 2007). The reference genome was annotated with the curated consensus sequences using RepeatMasker v. open-4.0.7 with a cut-off value of 250 (Smit, Hubley, and P. 2015). Simple repeats, low complexity regions and annotated elements shorter than 100 bp were filtered out and adjacent identical TEs overlapping by more than 100 bp were merged as belonging to the same TE family. Different TE families overlapping by more than 100 bp were considered as nested insertions and were renamed accordingly. Identical elements separated by less than 200 bp are indicative of interrupted elements and were grouped into a single element. TEs overlapping genes were recovered using the bedtools v. 2.27.1 suite and the “overlap” function (Quinlan and Hall 2010).

### POPULATION GENETIC ANALYSES

Genetic structure among isolates was estimated using unsupervised model-based Bayesian clustering implemented in STRUCTURE v. 2.34 as previously described (Mohd-Assaad, Mcdonald, and Croll 2018; Pritchard, Stephens, and Donnelly 2008). For principal component analysis (PCA) the “prcomp” function of the base R package stats was used. For linkage disequilibria decay estimations, the previously described SNP dataset was used and filtered for a minor allele frequency (MAF) > 5%. All calculations were made with VCFtools v. 0.1.12a (Danecek et al. 2011). Scans for selection were performed as described previously (Mohd-Assaad, Mcdonald, and Croll 2018). Briefly, integrated haplotype scores (iHS) and cross-population extended haplotype homozygosity (XP-EHH) were calculated as implemented in the R package rehh v. 2.02 (Gautier, Klassmann, and Vitalis 2017). The analysis was performed for each genetic cluster separately.

### GENOME WIDE ASSOCIATION STUDIES CNVS

We used the R package treeWAS to test for associations of CNV-alleles with phenotypic traits (Collins and Didelot 2018). TreeWAS was specifically developed for microbial population structures with significant degrees of clonality. Nine traits were tested (azole resistance, virulence, growth rate at 12 °C, 18 °C, 22 °C, temperature sensitivity and melanization at 12 °C, 18 °C, 22 °C). For each association analysis, the following parameters were used: tree = "BIONJ", n.subs = NULL, n.snps.sim = ncol(snps)*10, chunk.size = ncol(snps), test = c("terminal", "simultaneous", "subsequent"), snps.reconstruction = "parsimony", snps.sim.reconstruction = "parsimony", phen.reconstruction = "parsimony", na.rm = TRUE, p.value = 0.01, p.value.correct = "FDR", p.value.by = "count". P-values reported in the result are taken from the simultaneous reconstruction. The phylogenetic relationships between isolates were estimated by treeWAS.

#### GENOME WIDE ASSOCIATION STUDIES SNPS

We filtered the SNP data set retaining only SNPs with a genotyping rate >90% and a minor allelefrequency (MAF) >5%. GWAS analyses were performed as described previously (Mohd□ Assaad, McDonald, and Croll 2016). Briefly, we performed the analysis with TASSEL 20211027 using a mixed linear model (MLM) (Bradbury et al. 2007). To correct for the relatedness among individuals, we calculated a kinship matrix using the centered identity-by-state method (IBS) in TASSEL (Endelman and Jannink 2012). P-values were corrected using the base R “p.adjust” function.

## RESULTS

### IDENTIFICATION OF RECENT LARGE SEGMENTAL DUPLICATIONS AND SHORT DELETIONS

To assess the impact of CNV dynamics in *R. commune* populations across the worldwide geographic range of the pathogen, we first perform a thorough annotation of protein-encoding genes and repetitive sequences of the reference genome (Supplementary Table S1). To annotate coding sequences, we integrate RNA-sequencing evidence both in culture medium and across the infection cycle. The improved annotation includes 11’923 genes compared to 8’847 genes in previous annotations, which were based primarily on expressed sequence tags (ESTs; Supplementary Table S2 and S3) (Penselin et al. 2016). We functionally characterize all newly annotated gene models and additionally screen for candidate effector proteins and carbohydrate-active enzymes (CAZymes), both of which are known to be critical for host colonization and exploitation. To analyze repetitive sequences surrounding CNVs, we perform an exhaustive *de novo* annotation of TEs. In total, we classify 10’112 high-quality TE sequences grouped into 10 superfamilies.

We detect CNVs in 126 haploid *R. commune* isolates comparing read depth of Illumina sequencing data mapped to the reference genome (average depth of 37X) (Abyzov et al. 2011). We focus on polymorphic CNV events where both the gene presence and absence alleles are still segregating within the species. After quality filtering procedures (see Supplementary Fig. S1), we retain CNVs affecting a total of 1’861 distinct genes (15.6% of all genes). We find predominantly gene duplications (7’879 genes; 98.5%) and few gene deletions (116 genes or 1.5%; Fig. 1A). Duplications, but not deletions, often span across scaffolds, indicating large segmental duplications. In some isolates, multiple individual duplications amount to >200 kb each, suggesting that entire chromosomes may be duplicated. The few and small deletions indicate purifying selection against the deletion of essential genes contained in CNVs.

**Fig. 1.**
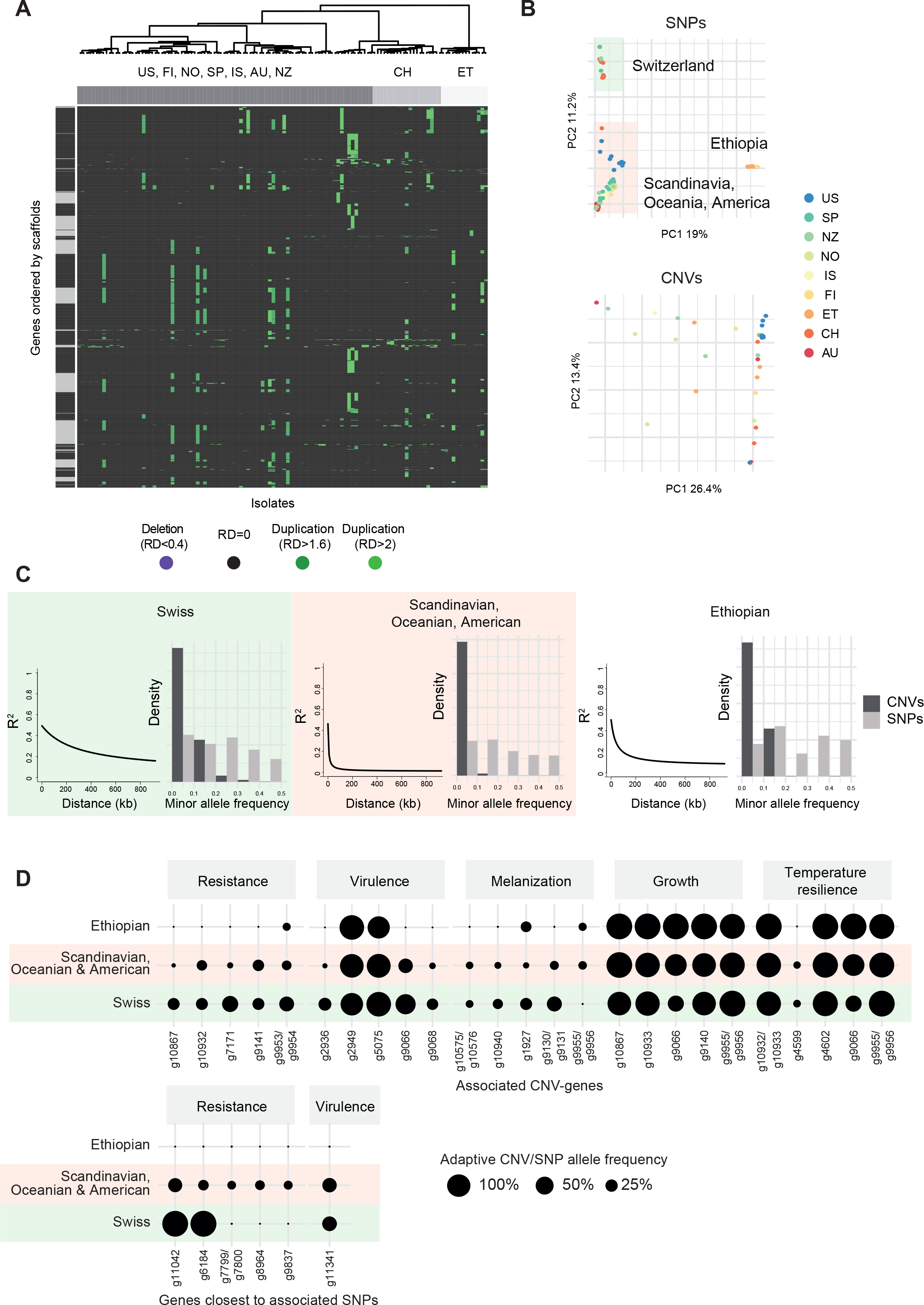
Segmental duplications and deletions co-segregate with genetic structure of *Rhynchosporium commune* populations. (A) Normalized read depth across genes and isolates. Read depth > 1.6 is indicated in green, read depth < 0.4 is indicated in purple. Genes affected by copy number variations (CNVs) are sorted by scaffolds (indicated as black and grey bars). Isolates are clustered based on single nucleotide polymorphisms (SNPs). (B) Principal component analysis based on SNPs (top) and based on CNVs in genes (bottom). Colors indicate country of origin: New Zealand (NZ), Australia (AU), Ethiopia (ET), Switzerland (CH), Spain (SP), Norway (NO), Finland (FI), Iceland (IS) and USA (US). (C) Decay of linkage disequilibrium and minor allele frequency spectra of CNVs within the three genetic clusters. Linkage disequilibrium between pairs of SNPs, measured as *r*^2^, is plotted against distance in kb. Minor allele frequency spectra are contrasted with the allele frequency of the derived allele at synonymous SNPs (n_CNVs_ and n_SNPs_ Swiss cluster = 780 and 71’842, Scandinavian, Oceanian & American cluster = 2061 and 219’357, Ethiopian cluster = 630 and 32’506). (D) Frequency of the trait associated CNV or SNP alleles identified with association analysis per genetic cluster. Alleles are associated with higher cypronazole resistance (measured as EC50), higher virulence (measured as PLACL), higher melanization at 18°C, higher growth at 18°C and higher temperature resilience (defined as standard deviation of the growth rate at 12°C, 18°C and 22°C). From all significantly associated CNV alleles with FDR-corrected p-values <0.05 only the top five are shown. For SNPs, no significant associations with melanization, growth or temperature resilience were found.

### POLYMORPHISM FREQUENCY SPECTRA ACROSS GENETIC CLUSTERS

To characterize the genetic differentiation across the pathogen distribution range, we perform an unsupervised Bayesian STRUCTURE analysis based on genome-wide single nucleotide polymorphisms (SNPs). We identify three well-supported clusters: a first cluster including isolates from Scandinavia, Oceania, and the United States, a second cluster separate from the other Northern European countries with mostly isolates from Switzerland and a third cluster with African isolates from Ethiopia. The clustering is consistent with a principal component analysis (Fig. 1B, upper panel). We analyze linkage disequilibrium decay within each cluster. The three clusters show very different rates of linkage disequilibrium (*r*^2^) decay indicating different degrees of clonality and demographic histories (Fig. 1C). The Scandinavian, Oceanian & North American cluster shows a very rapid linkage disequilibrium decay to *r*^2^=0.25 at an average distance of 2 kb, whereas the Swiss cluster shows the slowest decay to *r*^2^=0.25 at an average distance of 580 kb. Next, we analyze how selection acted on gene duplications and gene deletions in the different clusters. For this, we contrast the minor allele frequency spectrum of CNV alleles (*i.e.* duplication or deletion) and synonymous SNPs. We find a strong excess of low-frequency CNV alleles compared with synonymous SNPs in all clusters (Fig. 1C, Supplementary Fig. S2A, minor allele frequency ≤ 10%, Fisher’s exact test *p*- ≤ values < 0.001). In addition, we identify regions under positive selection using XP-EHH and iHS statistics and find that CNVs are depleted in positively selected regions (Supplementary Fig. S2B and Supplementary Table S4). This pattern suggests that both gene duplications and deletions are under strong negative selection in all clusters. To elucidate the genetic structure of CNV polymorphisms, we perform a principal component analysis based on CNV genotypes. The first principal component separates isolates primarily based on the total number of CNVs, with the isolate AU118 showing the highest number of CNVs compared to the reference genome (Supplementary Fig. S2C and D). The PCA does not reveal distinct clusters and shares little similarity with the PCA based on genome-wide SNPs, indicating that the majority of the detected CNVs was recently acquired (Fig. 1B, lower panel).

### TRAIT-ASSOCIATED CNVS AND SNPS SEGREGATING WITHIN GENETIC CLUSTERS

We then investigate the association of CNVs with phenotypic trait variation of *R. commune* isolates. For this, we perform a genome-wide association analysis of CNV-gene loci and nine traits including azole resistance, pathogenesis-related traits such as virulence (lesion coverage on leaves) and colony melanization at different temperatures. Additionally, we test for associations with colony growth rates at different temperatures as a proxy for temperature resilience. In total, we identify 191 genic CNV loci associated with one or more traits (FDR corrected *p*-value < 0.05, Supplementary Table S5). To provide a genomic context to the CNV-based association analyses, we repeat the association mapping with genome-wide SNPs. The GWAS with SNPs reveals significant associations for azole resistance and virulence, but not for the melanization and temperature related traits (FDR corrected *p*-value < 0.05, Supplementary Table S6, Fig. 1D). Among the significant associations for azole resistance and virulence, we find no overlap between SNPs and CNVs. We find considerable variation in the frequencies of the adaptive alleles among the three genetic clusters identified through CNV and SNP association mapping. This suggests distinct evolutionary histories of adaptation to the host, fungicide resistance and temperature resilience (Fig. 1D). Adaptive SNP and CNV alleles for azole resistance are frequent in the Swiss and the Scandinavian, Oceanian and North American cluster, but absent in the Ethiopian cluster, which is consistent with the largely inexistant use of azoles in Africa (“Food and Agriculture Organization of the United Nations (FAO) Pesticides Use Database” 2021).

### LARGE SEGMENTAL DUPLICATIONS ARE ASSOCIATED WITH PATHOGENESIS AND TEMPERATURE-RELATED TRAITS

Next, we characterize the individual CNV loci associated with trait variation. The encoded gene functions are enriched in nucleic acid or cofactor binding activity, *e.g.* proteins with endo-, exo- or ribonuclease domains, or oxygen binding proteins from the cytochrome p450 family (Fig. 2A). Furthermore, associated genes are enriched in catalytic activity, for example proteins with hydrolase and transferase activity. We find that CNV-genes associated with virulence are more strongly regulated during infection compared to non-associated CNV-genes, supporting their likely role in pathogenesis (Fig. 2B and Supplementary Fig. S3 A).

**Fig. 2.**
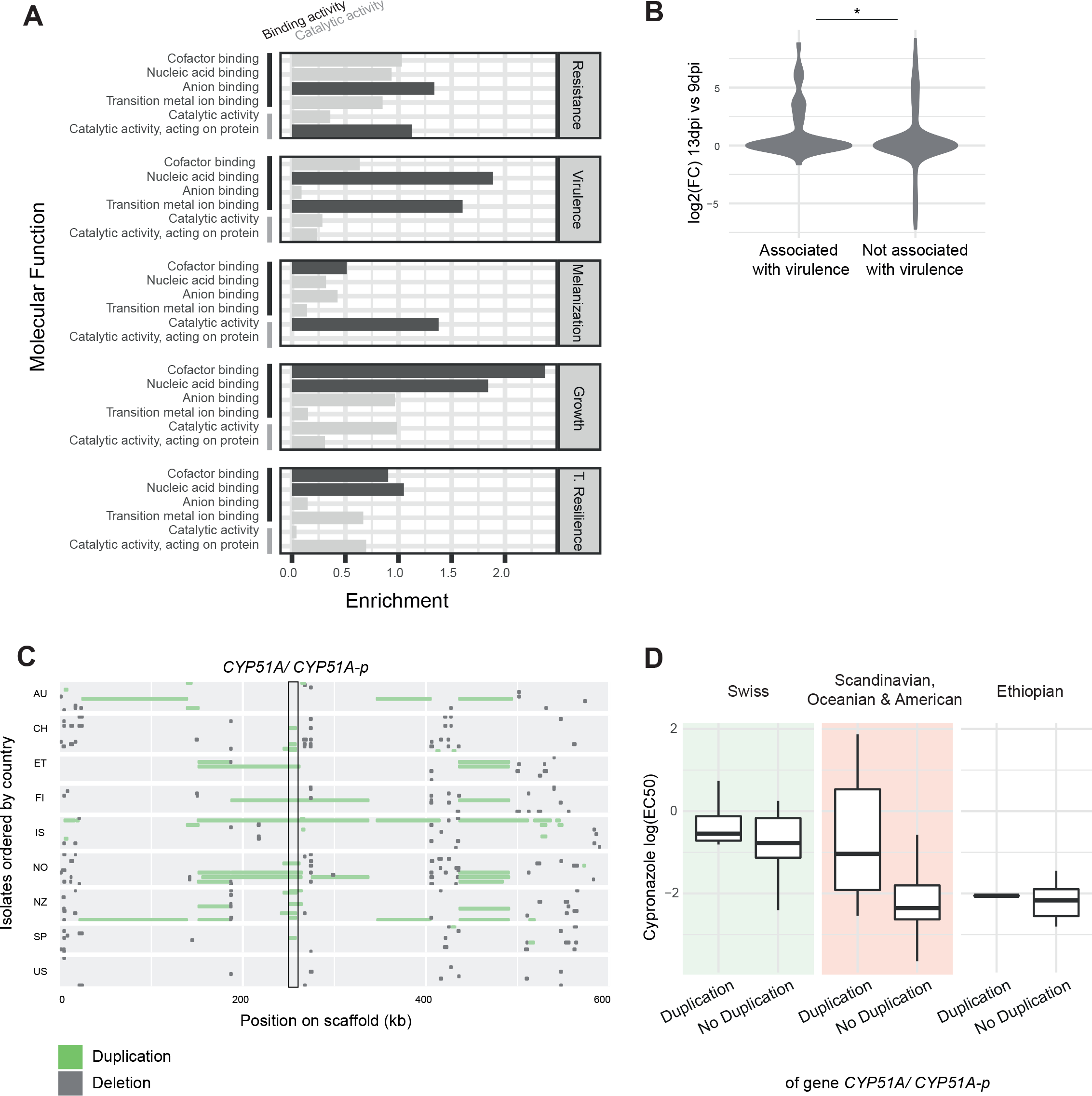
Associations of segmental duplications with azole resistance and virulence. (A) Molecular function gene ontology (GO) terms associated with cypronazole resistance (measured as EC50), virulence (measured as percent leaf area covered by lesions; PLACL), melanization at 18°C, growth at 18°C and temperature resilience (defined as the standard deviation among the growth rates at 12°C, 18°C and 22°C). The two most enriched categories for each trait are shown in dark grey. The enrichment score is defined as -log10(*p*-value). (B) Differential regulation upon infection for CNV-genes associated with virulence (genes with a *p*-value < 0.05, n = 53) and CNV-genes not associated with virulence (genes with a *p*-value > 0.05, n = 441). Differential regulation is defined as difference in RPKM between 13dpi and 9dpi. Virulence associated genes are significantly more upregulated than not associated genes (Wilcoxon-test p-value = 0.04). (C) Duplications (green) and deletions (grey) 200 kb up-and downstream of the cypronazole associated gene *CYP51A/ CYP51A-p* (marked with rectangle). The *CYP51A/ CYP51A-p* gene is spanned with many large segmental duplications. (D) Cypronazole resistance depicted as log(EC50) of *R.commune* isolates with and without duplication of the *CYP51A/ CYP51A-p* gene, grouped by genetic cluster (n_Duplication_ and n_No Duplication_ for the Swiss cluster = 4 and 10, Scandinavian, Oceanian & American cluster = 10 and 84, Ethiopian cluster = 1 and 12, *CYP51A/ CYP51A-p* association overall p-value < 0.0001).

Interestingly, the strongest association with azole resistance is the duplication of a sterol 14a- demethylase gene *CYP51A*/ *CYP51A-p* (g9954) and the adjacent gene encoding a calcineurin phosphoesterase (g9953) (Fig. 2C and Supplementary Fig. S3 B). The protein encoded by *CYP51* is the direct target of azole fungicides and overexpression and point mutations of *CYP51* have been shown to contribute to azole resistance in many crop pathogens (Lucas, Hawkins, and Fraaije 2015). Here, we observe that a large duplication of the region (2-200kb) harboring a *CYP51A* paralogue significantly contributes to variation in azole resistance (Fig. 2C). We find that the association between resistance and *CYP51A* paralogue duplication is strong in the Scandinavian, Oceanian and American cluster and the Switzerland cluster but not in the Ethiopian cluster (Fig. 2D). Only a single Ethiopian isolate carried a *CYP51A* duplication preventing a direct test for association. The low frequency in the Ethiopian cluster is likely due to a lack of azole application in this region compared to high azole usage in European or North American regions (“Food and Agriculture Organization of the United Nations (FAO) Pesticides Use Database” 2021). CNV calling based on short read sequences is not able to confidently distinguish between the *CYP51A* and *CYP51A-p* pseudogene paralogues due to their sequence similarity. Hence, the predicted duplication could indicate either a duplication of *CYP51A* or the pseudogene. Alternatively, indications for a duplication of *CYP51A* could also stem from a genotype carrying one copy of *CYP51A* and *CYP51A-p* each. Among the genes associated with virulence, 9.5% (*n* = 5) have homology to known effectors in other pathogens (Supplementary Table S4). Additionally, a CNV-gene associated to virulence overlaps with an open reading frame related to a retrotransposon from the *Copia* superfamily (g9066), another one with an epoxide hydrolase-like protein (g4694). Both genes show strong upregulation upon infection (Supplementary Table S3). Furthermore, a gene duplication with a strong virulence association (g9930) is located in a region showing signatures of a selective sweep (Supplementary Table S4). We also identify several genes encoding major facilitator superfamily (MFS) transporters with CNVs associated with different phenotypic traits. MFS transporters import and export sugars, metabolites and drugs and have known functions in both metabolism and pathogenicity in other pathogens (Costa et al. 2014; Del Sorbo, Schoonbeek, and De Waard 2000; Cavalheiro et al. 2018). A CNV of the gene encoding the MFS transporter g11393 is strongly associated with colony melanization across contrasted temperatures. For the gene encoding the MFS transporter g1792, we find an association with growth at low temperatures (12°C and 18°C). Taken together, CNV affecting genes likely contribute significantly to phenotypic trait variation across environments.

### CONSERVED AND ORPHAN GENE CNVS ARE ENRICHED IN DISTINCT ADAPTIVE FUNCTIONS

To investigate the evolutionary history of genes associated with CNVs in *R. commune*, we systematically analyze the presence of orthologues in other species. We define orthogroups across sister species, and 68 other Leotiomycetes species as well as four more distantly related Sordariomycetes (Supplementary Fig. S4). Overall, 27.2% of *R. commune* genes (n = 3243) share an orthologue in all Leotiomycetes and Sordariomycetes species, subsequently referred to as conserved genes. In contrast, 5.2% of *R. commune* genes (n= 623) have no orthologue in any other of the analyzed species, subsequently referred to as orphan genes (Fig. 3A and Supplementary Fig. S4). Out of 3,243 conserved genes, 671 (20.7%) are affected by CNV in *R. commune*, hence represent most likely a recent gene loss in the species. Conversely, 152 (24.4%) out of 623 orphan genes are affected by CNV in *R. commune*, hence the affected genes have most likely been gained since the last speciation (Fig. 3A). Gene loss alleles are overall at lower frequencies in the species compared to gene gain alleles, suggesting stronger purifying selection against recent gene losses (Fig. 3B).

**Fig. 3.**
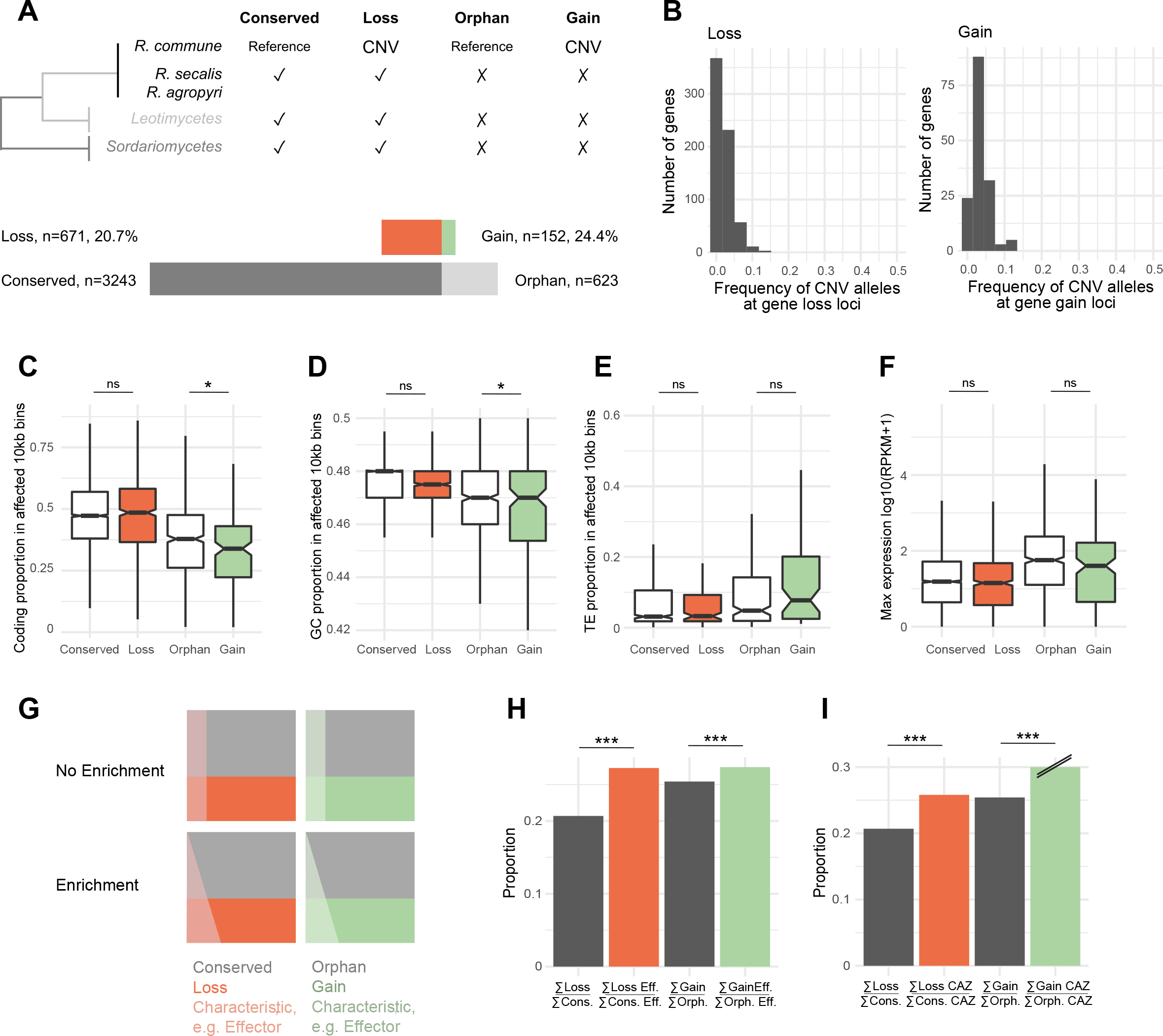
Adaptive functions of gain and copy-number variation (CNV). (A) Phylogenetic tree representing the relationships between *Rhynchosporium commune*, Leotimycetes and Sordariomycetes. Evidence for orthologues was used to infer whether a CNV in *R. commune* is likely due to a recent gene loss or gene gain. Numbers of genes for each category are indicated. (B) Frequency of CNVs at gene gain and gene loss loci for all isolates. (C)-(E) Differences between conserved, orphan, loss and gain genes regarding coding proportion (C), GC proportion (D) and transposable element proportion (E), each in proportion of affected 10 kb bins (two-sided t-test, * indicates p-values < 0.05). (F) Differences between conserved, orphan, loss and gain genes regarding maximal gene expression level during infection of a barley host, where RPKM is reads per kilobase of exon per million mapped reads. (G) Schematic representation of the enrichment analysis of adaptive gene functions. To infer if an adaptive gene group (i.e. effectors) is enriched in loss genes, the number of loss genes divided by conserved genes is compared to the fraction of the gene group in loss genes divided by the fraction of the gene group in conserved genes. Conversely, to infer if the gene group is enriched in gain genes, the number of gain genes divided by orphan genes is compared to the fraction of the gene group in gain genes divided by the fraction of the gene group in orphan genes. (H) Enrichment analysis for effector genes. The number of loss genes divided by the number of conserved genes is compared to the number of effector genes in loss genes divided by the number of effector genes in conserved genes. The number of gain genes divided by the number of orphan genes is compared to the number of effector genes in loss genes divided by the number of effector genes in orphan genes (Fisher-test loss-conserved p-value < 0.001, gain-orphan p-value < 0.001, n_Eff_ in conserved genes = 22, in loss genes = 6, in orphan genes = 73, in gain genes = 20). (I) Enrichment analysis for cell wall degrading enzymes (CAZymes). Proportions are calculated as above (Fisher-test loss-conserved p-value < 0.001, gain-orphan p-value < 0.001, n _CAZ_ in conserved genes = 62, in loss genes = 16, in orphan genes = 0, in gain genes = 1).

To characterize the genomic context of CNVs, we compare CNVs affecting conserved genes (*i.e.* gene losses) against all conserved genes. We find that loci with gene losses are in regions with similar coding density, GC and TE content, and show similar expression patterns as conserved genes (Fig. 3C-F and Supplementary Fig. S5). Similarly, we compare CNVs affecting orphan genes (*i.e.* gene gains) against all orphan genes. Recently gained genes are located in regions with lower coding density and GC content, but higher TE content than orphan genes in general. Recently gained genes also tend to show lower expression compared to orphan genes (Fig. 3C-F and Supplementary Fig. S5).

Subsequently, we investigate whether CNVs affecting conserved or orphan genes might impact host colonization. For this, we test whether CNV-genes are enriched in genes encoding candidate effector proteins and carbohydrate-active enzymes (CAZymes) (Fig. 3G). Effectors are small proteins secreted by the pathogen that manipulate the immune system and metabolism of the host to surmount plant resistance, whereas CAZymes are involved in the degradation of plant cell walls and nutrient acquisition (Esquerré-Tugayé, Boudart, and Dumas 2000; Langner and Göhre 2016; Fouché, Plissonneau, and Croll 2018; Sánchez-Vallet et al. 2018). To analyze CAZymes and effector candidates relevant for *R. commune* more comprehensively, we define orphan genes for this analysis at the genus level. We find that both genes associated with losses or gains are strongly enriched in effector candidates and CAZymes, suggesting that both gene losses and gene gains might contribute to host exploitation (Fig. 3H-I).

### TRANSPOSABLE ELEMENT ASSOCIATIONS WITH CNVS

Copy-number variation in genomes often arises through the presence of duplicated repetitive sequences(Hastings et al. 2009; Wells and Feschotte 2020). Overall, we find that almost half of the *R. commune* TEs belongs to the LTR retrotransposon *Gypsy* superfamily (RLG), followed by DNA-transposons with terminal inverted repeats (DTX), mostly members of the MITEs family, and LTR retrotransposons of the *Copia* superfamily (RLC) (Fig. 4A). To identify how repetitive elements might contribute to the emergence of CNVs in *R. commune*, we examine their distribution at CNV loci. We find that the *Gypsy* superfamily is underrepresented nearby CNVs. Instead, the most frequent TEs in proximity to CNV-genes are DNA-transposons from the *Tc1-Mariner* (DTT) and from the MITEs families, as well as from the *Copia* superfamily (Fig. 4B and Supplementary Fig. S6). Aditionally, the *Copia* superfamily is overrepresented in proximity to CNVs associated with phenotypic trait variation (Supplementary Fig. S6). In particular, we find that CNV-genes associated with azole resistance or virulence are closer to *Copia*, *Tc1-Mariner* and MITEs compared to CNV-genes with no association with trait variation (Fig. 4C and Supplementary Fig. S6). Among the CNV- genes with an association to virulence, we find a gene encoding a metallopeptidase with an overlapping MITE, as well as an endonuclease gene with an overlapping *Tc1-Mariner* element. Both insertions have possibly caused gene disruptions (Fig. 4D and Supplementary Fig. S7 A). The metallopeptidase protein familiy has been shown to play a role in virulence in other plant pathogens (Yike 2011). Furthermore, several of the azole resistance associated genes encode for open reading frames of *Copia* elements (Fig. 4E and Supplementary Fig. S7 A). To further investigate the potential role of MITEs, *Tc1-Mariner* and *Copia* TEs in *R. commune* adaptation, we test whether copies of these elements are enriched in regions under positive selection. We find that copies of MITEs are strongly enriched in regions with signatures of recent positive selection compared to the genomic background. On the contrary, copies of *Gypsy* elements are strongly depleted in regions with signatures of recent positive selection (Fig. 4F and Supplementary Table S4). The enrichment and depletion signatures are consistent across the three genetic clusters (Supplementary Fig. S7 B). Moreover, we find that copies of MITEs have the highest GC content among all superfamilies (Fig. 4G). MITEs are also strongly up-regulated upon infection, consistent with a stress-induced de-repression of these elements (Fig. 4H). Taken together, MITEs are likely playing a key role in generating recent adaptive CNVs within the species.

**Fig. 4.**
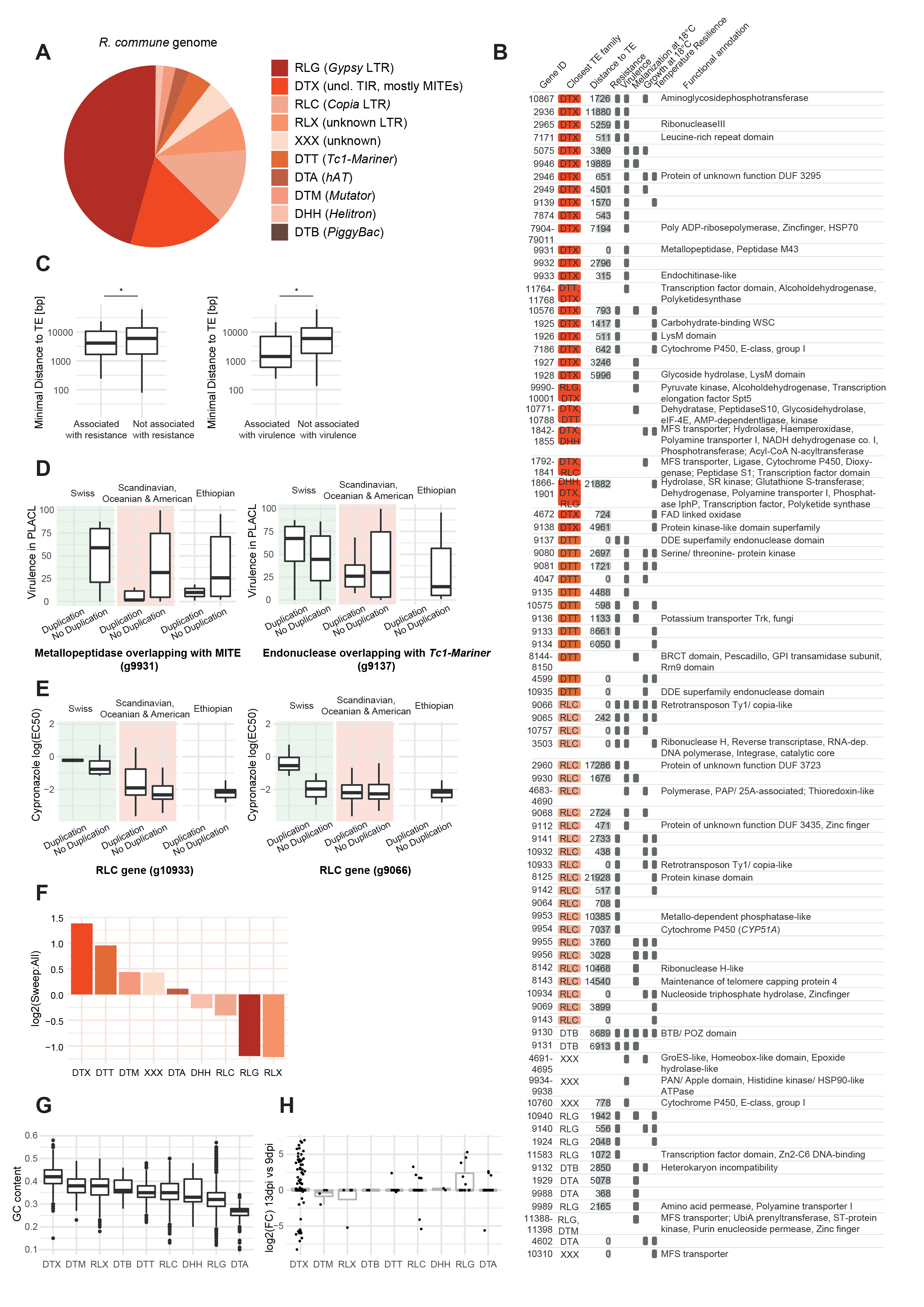
Roles of MITEs, *Tc1-Mariner* and *Copia* transposable element (TE) families in adaptation. (A) TE superfamily distribution in the *R.commune* genome, specifically DNA-TEs of the superfmailies *hAT* (DTA), *PiggyBac* (DTB), *Tc1-Mariner* (DTT), *Helitron* (DHH), *Mutator* (DTM) and unclassified TIR (DTX), as well as LTR retrotransposon superfamilies *Copia* (RLC), *Gypsy* (RLG) and unclassified LTR (RLX), as well as unclassified TEs (XXX). (B) Gene functions and distance to the closest TE are shown for CNV-genes associated with cypronazole resistance (measured as EC50), virulence (measured as PLACL), melanization at 18°C, growth at 18°C and temperature resilience (defined as standard deviation of the growth rate at 12°C, 18°C and 22°C). Shown are CNV-genes with an association of p-value < 0.01 and the associated traits are highlighted. Gene descriptions are based on Interpro annotations. For each gene, the distance to the closest TE and the respective TE superfamily is indicated. Genes are ordered by TE superfamily. (C) Minimal distance to TEs of CNV-genes that are associated with resistance (left) and virulence (right), compared to not associated CNV-genes. Associated CNV-genes are closer to TEs than not associated genes (Wilcoxon-test p-value = 0.03 and < 0.001, n_Associated_ and n_Not Associated_ resistance n = 34 and n = 325, and virulence n = 26 and n = 333). Association is defined as p-value < 0.05, non-associated as p-value > 0.05 respectively. Only DTX, DTT and RLC TE superfamilies are included. (D) Examples of virulence associated CNV-genes close to DTX and DTT TEs. Virulence is depicted as PLACL of *R.commune* isolates with and without gene duplication, grouped by genetic cluster (g9931: n_Duplication_ and n_No Duplication_ for the Swiss cluster = 0 and 12, Scandinavian, Oceanian & American cluster = 5 and 85, Ethiopian cluster = 2 and 10; g9137: n_Duplication_ and n_No Duplication_ for the Swiss cluster = 4 and 8, Scandinavian, Oceanian & American cluster = 5 and 85, Ethiopian cluster = 0 and 12, p-values of associations < 0.05). (E) Examples of cypronazole resistance associated CNV-genes close to RLC TEs. Cypronazole resistance is depicted as log(EC50) of *R.commune* isolates with and without gene duplication, grouped by genetic cluster (g10933: n_Duplication_ and n_No Duplication_ for the Swiss cluster = 2 and 12, Scandinavian, Oceanian & American cluster = 10 and 84, Ethiopian cluster = 0 and 13; g9066: n_Duplication_ and n_No Duplication_ for the Swiss cluster = 12 and 2, Scandinavian, Oceanian & American cluster = 28 and 66, Ethiopian cluster = 0 and 13, p-values of associations < 0.05). (F) TE superfamily distribution in *R. commune* sweep regions identified by XP-EHH and iHS. Bar diagram indicates over- respectively underrepresentation of TE superfamilies in sweep regions compared to the whole genome. DTX TEs are the most enriched in sweep regions, whereas RLG are the most depleted (Fisher’s exact test DTX and RLG p-values < 0.001). (G) GC content of TE superfamilies. DTX TEs show the highest GC content. (H) Differential regulation upon infection of genes overlapping with different TE superfamilies. Differential regulation is defined as difference in RPKM between 13dpi and 9dpi. Genes overlapping with DTX TEs show strong differential expression.

## DISCUSSION

Pathogens in agricultural environments are under strong selection pressure to adapt to changing environments. CNVs have been increasingly recognized as important raw material for adaptive evolution. We identify a substantial set of CNV loci associated with phenotypic trait variation across genetically differentiated global populations of the barley pathogen *R. commune*. Genome-wide association analyses based on SNPs shows contrasts in the number and frequencies of adaptive alleles compared to CNVs. We show that adaptation through CNVs was likely facilitated by the action of a specific set of TE families.

We use a broad phylogenomic framework of Sordariomycetes and Leotiomycetes fungi to distinguish gene gain and loss events triggered by CNVs. We identify more recent gene losses than gene gains, however purifying selection acts more strongly against gene losses. Loci experiencing gene losses tend to encode essential protein functions. As our framework for gene loss identification is based on conservation across a broad phylogeny, such a tendency is in part due to our reliance on phylogenetic signals. Recent gene gains are selectively mostly neutral. This matches expectations for young genes often lacking essential functions at early stages of gain-of-function processes (McLysaght and Guerzoni 2015). Interestingly, both, gene losses and gains are enriched in functions related to host interactions including effector candidates, which may enable the pathogen to manipulate the host physiology. Gains and losses are also enriched in genes encoding CAZymes involved in the degradation of plant cell walls and nutrient acquisition (Esquerré-Tugayé, Boudart, and Dumas 2000; Langner and Göhre 2016). The increased gene gain and loss rates for functions related to host exploitation may be driven by strong selection on pathogen populations to gain new functions and evade detection of proteins exposed to the host immune system. Consistent with the over- representation of host exploitation functions at CNV loci, we find strong support for the role of individual CNVs in association with pathogenesis-related trait variation. We find that pathogenesis associated CNV-genes are enriched for catalytic activity and cofactor binding activity. Catalytic activity is consistent with the molecular functions of CAZymes, whereas cofactor binding activity is consistent with effector associated functions (Lo Presti et al. 2015; Esquerré-Tugayé, Boudart, and Dumas 2000). Gene deletions at effector loci have previously been shown to underlie substantial variation in pathogenicity within crop pathogen species (Hartmann et al. 2017; Lo Presti et al. 2015). In *Z. tritici*, a segregating gene deletion largely explains the gain of virulence on a cognate wheat cultivar (Hartmann et al. 2017). Our CNV association mapping does not detect relevant associations at the *NIP* effector gene loci, however this is largely explained by the fact that the tested barley cultivar lacks the corresponding receptor (Mohd□ Assaad, McDonald, and Croll 2019). The association mapping also reveals a series of associations with temperature tolerance and fungicide resistance traits, showing how extensive the species segregates adaptive genetic variation at the level of gene presence/absence polymorphism.

We find that most CNVs underlying phenotypic trait variation are large segmental duplications affecting multiple genes. The most striking duplication associated with fungicide resistance includes the *CYP51A* gene known to encode the target of azoles. Using targeted PCR assays, previous studies have found a correlation between azole sensitivity and the presence of the *CYP51A* paralogue in *R. commune* (Hawkins et al. 2014; Brunner et al. 2016). Here, we find that CNV variation at the *CYP51A/CYP51A-p* loci is relevant for azole resistance in a global population genetic context. However, the short read mapping approach for CNV calling cannot confidently distinguish between the close paralogues *CYP51A* and *CYP51A-p* due to the high degree of sequence similarity. Hence, we cannot pinpoint which of the two paralogues is causal for the variation in resistance. However, based on previous analysis, the *CYP51A-p* contains premature stop codons and mutations do not contribute to azole resistance, therefore we assume that at least part of the CNV variation we observe is related to *CYP51A* (Hawkins et al. 2014; Brunner et al. 2016). In human pathogenic yeasts such as *C. neoformans* and *C. albicans,* several chromosomal duplications have been associated with *ERG11*, the *CYP51* orthologue (Coste et al. 2007; Sionov et al. 2010). The amplification of the locus is typically observed over the course of a single infection under strong selection pressure. Chromosomal aberrations are often accompanied with significant fitness reductions and lost in absence of fungicide use (Zhang et al. 2019; Morschhäuser 2016). In plant pathogens, complete chromosomal duplications of *CYP51* have not been observed so far. However, many ascomycetes show several deeply divergent *CYP51* paralogues, whereas *C. albicans* and *C. neoformans* have only a single copy of *CYP51* (Zhang et al. 2019). In *R. commune*, we find that the duplication affecting *CYP51A* ranges from 2-200kb in different isolates. Hence, it seems plausible that in some *R. commune* isolates, the *CYP51A* amplification arose analogously as in human pathogens from partial chromosomal duplications. Although we cannot decipher the exact mechanisms that led to the *CYP51A* duplication in *R. commune* observed in this study, the large chromosomal duplications share similarities to the process of intrahost fungicide resistance evolution in human pathogens (Coste et al. 2007; Sionov et al. 2010). Further studies will be needed to identify the exact mechanisms triggering chromosomal duplications in human and predominantly clonal plant pathogens such as *R. commune*.

Adaptive genetic variation is typically mapped using genome-wide SNPs because such markers can densely cover most chromosomal regions. At high marker density, SNPs tend to associate with additional types of polymorphisms (*e.g.* insertions, deletions and duplications) segregating in populations through linkage disequilibrium (Tam et al. 2019). Hence, association mapping performed on SNPs is expected to unravel associated polymorphisms even if the causal variant is not a SNP. Here, we analyze the outcomes of SNP- and CNV-based association mapping and find unexpectedly large discrepancies. Analyzing SNPs, we identify associations with fungicide resistance, but not with temperature related traits. The dominant role of CNVs compared to SNPs in temperature adaptation has been identified previously (Dorant et al. 2020). We identify no gene associated both in SNP- and CNV-based association mapping. The two sets of markers reveal also unequal numbers of associations among genetic clusters and differences in adaptive allele frequencies at significantly associated loci. Overall, our comparison of SNP- and CNV-based association mapping shows how including different types of polymorphisms improves the coverage of genome-wide genetic variation. Our findings also affirm that a comprehensive analysis of polymorphisms is necessary to unravel the genetic architecture of complex traits.

Complex sequence rearrangements have been repeatedly associated with specific TEs inducing non- homologues recombination events (Wells and Feschotte 2020; Bourque et al. 2018). Specifically in absence of meiosis, TE-mediated recombination could provide an important source of adaptive variability. In the partially clonal pathogen *R. commune,* we find that *Gypsy* retrotransposons make up about half of the TE copies in the genome, similar to many other fungal pathogens including *Blumeria graminis* and *Zymoseptoria tritici*, where retrotransposons contributed to recent genome expansions (Frantzeskakis et al. 2018; Oggenfuss et al. 2021). Next to *Gypsy* elements, *R. commune* shows a high frequency of DNA transposons with terminal inverted repeats (TIR), mostly from the MITES and *Tc1-Mariner* families. As previously observed, we find that MITES and *Copia* elements are located preferentially close to genes (Casacuberta et al. 1998; Santiago et al. 2002; Oki et al. 2008; Castanera et al. 2021). Interestingly, *Copia* elements are overrepresented nearby genes affected by CNVs. The overrepresentation is even more pronounced nearby CNV-genes associated with phenotypic trait variation. In plants, adaptive *Copia* element insertions are frequently observed, and the elements often have regulatory functions (Kanazawa et al. 2009; White, Habera, and Susan 1994; Baduel et al. 2021). In plant pathogens, retrotransposon elements have been found to be de-repressed upon stress with consequences for virulence on the host (Dubin, Mittelsten Scheid, and Becker 2018; Fouché et al. 2020). For example, in *Z. tritici*, *Copia* elements are upregulated during early infection, whereas other TE elements are only upregulated during the less stressful saprophytic stages of the pathogen life cycle (Fouché et al. 2020). CNV-genes associated with resistance or virulence are more closely located to *Copia*, MITEs and *Tc1-Mariner* elements than non-associated CNV-genes. The retention of MITEs in proximity to genes likely stems from the comparatively minor impact of short element insertions near coding sequences under purifying selection. However, the retention of some TEs nearby genes likely reflects also positive selection favoring *e.g.* an adaptive regulatory variant triggered by the insertion event (Castanera et al. 2021). MITEs are indeed known to have expression patterns that aid the infection process in several pathogens (Schmidt et al. 2013). It has been previously shown that MITEs allow to predict novel virulence genes in *Fusarium oxysporum* (Schmidt et al. 2013). In line with this, we find that MITEs are strongly regulated upon host infection.

Furthermore, we find that MITEs are enriched in selective sweep regions. Together, our results suggest an involvement of specific TE families in adaptive CNV generation. Although we cannot make the direct link between TE proliferation and CNV here, our results suggest that specific TE activity is connected to adaptive trait variation.

Our analyses of populations spanning the global distribution range of a crop pathogen allows us to track trait associated CNV-genes across geographic regions. We find that trait associated CNVs in close proximity to specific TEs show high degrees of variation in adaptive allele frequencies. Such high degree of frequency variation among genetic clusters possible reflects the recency of the onset of selection, the time since the CNV has first appeared in the cluster, or local adaptation. In particular, we find that CNV-genes associated with resistance to fungicides show higher allele frequency variation among clusters compared to CNV-genes associated with temperature adaptation, which are often close to fixation in all populations. This is consistent with the heterogeneous and time-shifted application of fungicides across the globe (“Food and Agriculture Organization of the United Nations (FAO) Pesticides Use Database” 2021). Understanding the population genetic context of microbial adaptation in agricultural ecosystems across the globe is a critical foundation to address risks of pathogen spread. Locally adapted pathogen populations may not benefit from gene flow to different regions in the world and may be more easily contained (Croll and McDonald 2017). However, homogenization of global agricultural ecosystems at the level of applied fungicides or deployed host genotypes reduces the constraints on locally adapted pathogen populations (Croll and McDonald 2017). Local adaptation may under such circumstances even favor gene flow to locations sharing similar host or abiotic environments. A mechanistic understanding of microbial adaptation furthermore improves our ability to predict the speed at which pathogen populations can cope with environmental challenges. Adaptation mediated by CNVs generated in repetitive regions of the genome can progress particularly fast (Mérot et al. 2020). Deprioritizing easily surmountable abiotic or biotic pressures on pathogen populations can improve the sustainability of pathogen control.

## Author contributions

LS and DC conceived the study and designed analyses, LS performed analyses, UO annotated and classified TEs, NMA provided samples/datasets, DC provided funding, LS and DC wrote the manuscript with input from co-authors. All authors reviewed the manuscript and agreed on submission.

## Data availability

The genome sequencing data for *Rhynchosporium commune* is available at the NCBI Sequence Read Archive under BioProject accession number PRJNA327656. The RNA sequencing data is available at https://doi.org/10.5281/zenodo.5729968 and https://doi.org/10.5281/zenodo.5729863, the updated genome annotation is available at https://doi.org/10.5281/zenodo.5730007.

## Acknowledgements

We thank Nikhil Singh and Thomas Badet for helpful discussions and comments on previous versions of the manuscript.

[dataset] Mohd-Assaad Norfarhan, Croll Daniel; 2016; Population genomics of Rhynchosporium commune; NCBI Sequence Read Archive; PRJNA327656

[dataset] Stalder Luzia, Oggenfuss Ursula, Mohd-Assaad Norfarhan, Croll Daniel; 2021; Illumina TruSeq stranded mRNA sequences of Rhynchosporium commune isolate UK7 in plantae; 10.5281/zenodo.5729968

[dataset] Stalder Luzia, Oggenfuss Ursula, Mohd-Assaad Norfarhan, Croll Daniel; 2021; Improved genome annotation of Rhynchosporium commune isolate UK7 using Illumina short reads of in vitro and in plantae conditions; 10.5281/zenodo.5730007

[dataset] Stalder Luzia, Oggenfuss Ursula, Mohd-Assaad Norfarhan, Croll Daniel; 2021; Illumina TruSeq stranded mRNA sequences of Rhynchosporium commune isolate UK7 in vitro; 10.5281/zenodo.5729863

## References

Abyzov, Alexej, Alexander E. Urban, Michael Snyder, and Mark Gerstein. 2011. “CNVnator: An Approach to Discover, Genotype, and Characterize Typical and Atypical CNVs from Family and Population Genome Sequencing.” Genome Research 21 (6): 974–84. https://doi.org/10.1101/gr.114876.110.

Altschul, Stephen F., Thomas L. Madden, Alejandro A. Schäffer, Jinghui Zhang, Zheng Zhang, Webb Miller, and David J. Lipman. 1997. “Gapped BLAST and PSI-BLAST: A New Generation of Protein Database Search Programs.” Nucleic Acids Research 25 (17): 3389–3402. https://doi.org/10.1016/B978-1-4832-3211-9.50009-7.

Anders, Simon, Paul Theodor Pyl, and Wolfgang Huber. 2015. “HTSeq-A Python Framework to Work with High-Throughput Sequencing Data.” Bioinformatics 31 (2): 166–69. https://doi.org/10.1093/bioinformatics/btu638.

Azevedo, Maria Manuel, Isabel Faria-Ramos, Luísa Costa Cruz, Cidália Pina-Vaz, and Acácio Gonçalves Rodrigues. 2015. “Genesis of Azole Antifungal Resistance from Agriculture to Clinical Settings.” Journal of Agricultural and Food Chemistry 63 (34): 7463–68. https://doi.org/10.1021/acs.jafc.5b02728.

Baduel, Pierre, Basile Leduque, Amandine Ignace, Isabelle Gy, José Gil, Olivier Loudet, Colot Vincent, et al. 2021. “Genetic and Environmental Modulation of Transposition Shapes the Evolutionary Potential of Arabidopsis Thaliana.” Genome Biology 22.

Bao, Weidong, Kenji K. Kojima, and Oleksiy Kohany. 2015. “Repbase Update, a Database of Repetitive Elements in Eukaryotic Genomes.” Mobile DNA 6 (1): 4–9. https://doi.org/10.1186/s13100-015-0041-9.

Bolger, Anthony M., Marc Lohse, and Bjoern Usadel. 2014. “Trimmomatic: A Flexible Trimmer for Illumina Sequence Data.” Bioinformatics 30 (15): 2114–20. https://doi.org/10.1093/bioinformatics/btu170.

Bourque, Guillaume, Kathleen H. Burns, Mary Gehring, Vera Gorbunova, Andrei Seluanov, Molly Hammell, Michaël Imbeault, et al. 2018. “Ten Things You Should Know about Transposable Elements.” Genome Biology 19 (199). https://doi.org/10.1186/s13059-018-1577-z.

Bradbury, Peter J., Zhiwu Zhang, Dallas E. Kroon, Terry M. Casstevens, Yogesh Ramdoss, and Edward S. Buckler. 2007. “TASSEL: Software for Association Mapping of Complex Traits in Diverse Samples.” Bioinformatics 23 (19): 2633–35. https://doi.org/10.1093/bioinformatics/btm308.

Breen, James, Thomas Wicker, Xiuying Kong, Juncheng Zhang, Wujun Ma, Etienne Paux, Catherine Feuillet, Rudi Appels, and Matthew Bellgard. 2010. “A Highly Conserved Gene Island of Three Genes on Chromosome 3B of Hexaploid Wheat: Diverse Gene Function and Genomic Structure Maintained in a Tightly Linked Block.” BMC Plant Biology 10. https://doi.org/10.1186/1471-2229-10-98.

Brown, Gordon D., David W. Denning, Neil A.R. Gow, Stuart M. Levitz, Mihai G. Netea, and Theodore C. White. 2012. “Hidden Killers: Human Fungal Infections.” Science Translational Medicine 4 (165). https://doi.org/10.1126/scitranslmed.3004404.

Brunner, Patrick C., Tryggvi S. Stefansson, James Fountaine, Veronica Richina, and Bruce A. McDonald. 2016. “A Global Analysis of CYP51 Diversity and Azole Sensitivity in Rhynchosporium Commune.” Phytopathology 106 (4): 355–61. https://doi.org/10.1094/PHYTO-07-15-0158-R.

Casacuberta, Elena, Josep M. Casacuberta, Pere Puigdomènech, and Amparo Monfort. 1998. “Presence of Miniature Inverted-Repeat Transposable Elements (MITEs) in the Genome of Arabidopsis Thaliana: Characterisation of the Emigrant Family of Elements.” Plant Journal 16 (1): 79–85. https://doi.org/10.1046/j.1365-313X.1998.00267.x.

Castanera, Raul, Pol Vendrell-Mir, Amelie Bardil, Marie Christine Carpentier, Olivier Panaud, and Josep Casacuberta. 2021. “Amplification Dynamics of Miniature Inverted-Repeat Transposable Elements and Their Impact on Rice Trait Variability.” The Plant Journal, no. 107: 118–35. http://biorxiv.org/cgi/content/short/2020.10.01.322784v1?rss=1&utm_source=researcher_app&utm_medium=referral&utm_campaign=RESR_MRKT_Researcher_inbound.

Cavalheiro, Mafalda, Pedro Pais, Mónica Galocha, and Miguel C. Teixeira. 2018. “Host-Pathogen Interactions Mediated by MDR Transporters in Fungi: As Pleiotropic as It Gets!” Genes MDPI 9 (332). https://doi.org/10.3390/genes9070332.

Collins, Caitlin, and Xavier Didelot. 2018. “A Phylogenetic Method to Perform Genome-Wide Association Studies in Microbes That Accounts for Population Structure and Recombination.” PLoS Computational Biology 14 (2). https://doi.org/10.1371/journal.pcbi.1005958.

Cools, Hans J., and Bart A. Fraaije. 2013. “Update on Mechanisms of Azole Resistance in Mycosphaerella Graminicola and Implications for Future Control.” Pest Management Science 69 (2): 150–55. https://doi.org/10.1002/ps.3348.

Costa, Catarina, Paulo J. Dias, Isabel Sá-Correia, and Miguel C. Teixeira. 2014. “MFS Multidrug Transporters in Pathogenic Fungi: Do They Have Real Clinical Impact?” Frontiers in Physiology 5. https://doi.org/10.3389/fphys.2014.00197.

Coste, Alix, Anna Selmecki, Anja Forche, Dorothée Diogo, Marie Elisabeth Bougnoux, Christophe D’Enfert, Judith Berman, and Dominique Sanglard. 2007. “Genotypic Evolution of Azole Resistance Mechanisms in Sequential Candida Albicans Isolates.” Eukaryotic Cell 6 (10): 1889– 1904. https://doi.org/10.1128/EC.00151-07.

Croll, Daniel, and Bruce A. McDonald. 2017. “The Genetic Basis of Local Adaptation for Pathogenic Fungi in Agricultural Ecosystems.” Molecular Ecology 26 (7): 2027–40. https://doi.org/10.1111/mec.13870.

Danecek, Petr, Adam Auton, Goncalo Abecasis, Cornelis A. Albers, Eric Banks, Mark A. DePristo, Robert E. Handsaker, et al. 2011. “The Variant Call Format and VCFtools.” Bioinformatics 27 (15): 2156–58. https://doi.org/10.1093/bioinformatics/btr330.

Dorant, Yann, Hugo Cayuela, Kyle Wellband, Martin Laporte, Quentin Rougemont, Claire Mérot, Eric Normandeau, Rémy Rochette, and Louis Bernatchez. 2020. “Copy Number Variants Outperform SNPs to Reveal Genotype–Temperature Association in a Marine Species.” Molecular Ecology 29 (24): 4765–82. https://doi.org/10.1111/mec.15565.

Dubin, Manu J., Ortrun Mittelsten Scheid, and Claude Becker. 2018. “Transposons: A Blessing Curse.” Current Opinion in Plant Biology 42: 23–29. https://doi.org/10.1016/j.pbi.2018.01.003.

Emms, David M., and Steven Kelly. 2019. “OrthoFinder: Phylogenetic Orthology Inference for Comparative Genomics.” Genome Biology 20 (238). https://doi.org/10.1101/466201.

Endelman, Jeffrey B., and Jean Luc Jannink. 2012. “Shrinkage Estimation of the Realized Relationship Matrix.” G3: Genes, Genomes, Genetics 2 (11): 1405–13. https://doi.org/10.1534/g3.112.004259.

Esquerré-Tugayé, Marie Thérèse, Georges Boudart, and Bernard Dumas. 2000. “Cell Wall Degrading Enzymes, Inhibitory Proteins, and Oligosaccharides Participate in the Molecular Dialogue between Plants and Pathogens.” Plant Physiology and Biochemistry 38 (1–2): 157–63. https://doi.org/10.1016/S0981-9428(00)00161-3.

Feschotte, Cédric. 2008. “Transposable Elements and the Evolution of Regulatory Networks.” Nature Reviews. Genetics 9 (May): 397–405.

Fisher, Matthew C., Daniel A. Henk, Cheryl J. Briggs, John S. Brownstein, Lawrence C. Madoff, Sarah L. McCraw, and Sarah J. Gurr. 2012. “Emerging Fungal Threats to Animal, Plant and Ecosystem Health.” Nature 484 (7393): 186–94. https://doi.org/10.1038/nature10947.

Fisher, Matthew C, Nichola J Hawkins, Dominique Sanglard, and Sarah J Gurr. 2018 Worldwide Emergence of Resistance to Antifungal Drugs Challenges Human Health and Food Security.” Science (New York, N.Y.) 360: 739–42. http://tclocal.org/2011/01/health_and_food_security.html.

Fones, Helen N., Daniel P. Bebber, Thomas M. Chaloner, William T. Kay, Gero Steinberg, and Sarah J. Gurr. 2020. “Threats to Global Food Security from Emerging Fungal and Oomycete Crop Pathogens.” Nature Food 1 (6): 332–42. https://doi.org/10.1038/s43016-020-0075-0.

“Food and Agriculture Organization of the United Nations (FAO) Pesticides Use Database.” 2021. http://www.fao.org/faostat/. 2021.

Fouché, Simone, Thomas Badet, Ursula Oggenfuss, Clémence Plissonneau, Carolina Sardinha Francisco, and Daniel Croll. 2020. “Stress-Driven Transposable Element De-Repression Dynamics and Virulence Evolution in a Fungal Pathogen.” Molecular Biology and Evolution 37 (1): 221–39. https://doi.org/10.1093/molbev/msz216.

Fouché, Simone, Clémence Plissonneau, and Daniel Croll. 2018. “The Birth and Death of Effectors in Rapidly Evolving Filamentous Pathogen Genomes.” Current Opinion in Microbiology 46: 34– 42. https://doi.org/10.1016/j.mib.2018.01.020.

Frantzeskakis, Lamprinos, Barbara Kracher, Stefan Kusch, Makoto Yoshikawa-Maekawa, Saskia Bauer, Carsten Pedersen, Pietro D. Spanu, Takaki Maekawa, Paul Schulze-Lefert, and Ralph Panstruga. 2018. “Signatures of Host Specialization and a Recent Transposable Element Burst in the Dynamic One-Speed Genome of the Fungal Barley Powdery Mildew Pathogen.” BMC Genomics 19 (381). https://doi.org/10.1186/s12864-018-4750-6.

Gautier, Mathieu, Alexander Klassmann, and Renaud Vitalis. 2017. “Rehh 2.0: A Reimplementation of the R Package Rehh to Detect Positive Selection from Haplotype Structure.” Molecular Ecology Resources 17 (1): 78–90. https://doi.org/10.1111/1755-0998.12634.

Hartmann, Fanny E., and Daniel Croll. 2017. “Distinct Trajectories of Massive Recent Gene Gains and Losses in Populations of a Microbial Eukaryotic Pathogen.” Molecular Biology and Evolution 34 (11): 2808–22. https://doi.org/10.1093/molbev/msx208.

Hartmann, Fanny E., Andrea Sánchez-Vallet, Bruce A. McDonald, and Daniel Croll. 2017. “A Fungal Wheat Pathogen Evolved Host Specialization by Extensive Chromosomal Rearrangements.” ISME Journal 11 (5): 1189–1204. https://doi.org/10.1038/ismej.2016.196.

Hastings, P. J., James R. Lupski, Susan M. Rosenberg, and Grzegorz Ira. 2009. “Mechanisms of Change in Gene Copy Number.” Nature Reviews Genetics 10 (8): 551–64. https://doi.org/10.1038/nrg2593.

Hawkins, J. Nichola, Hans J. Cools, Helge Sierotzki, Michael W. Shaw, Wolfgang Knogge, Steven L. Kelly, Diane E. Kelly, and Bart A. Fraaije. 2014. “Paralog Re-Emergence: A Novel, Historically Contingent Mechanism in the Evolution of Antimicrobial Resistance.” Mol. Biol. Evol. 31 (7): 1793–1802. https://pubmed.ncbi.nlm.nih.gov/24732957/?from_term=rhynchosporium+commune&from_page=2&from_pos=10.

Hoff, Katharina J., Simone Lange, Alexandre Lomsadze, Mark Borodovsky, and Mario Stanke. 2016. “BRAKER1: Unsupervised RNA-Seq-Based Genome Annotation with GeneMark-ET and AUGUSTUS.” Bioinformatics 32 (5): 767–69. https://doi.org/10.1093/bioinformatics/btv661.

Jones, Philip, David Binns, Hsin Yu Chang, Matthew Fraser, Weizhong Li, Craig McAnulla, Hamish McWilliam, et al. 2014. “InterProScan 5: Genome-Scale Protein Function Classification.” Bioinformatics 30 (9): 1236–40. https://doi.org/10.1093/bioinformatics/btu031.

Kanazawa, Akira, Baohui Liu, Fanjiang Kong, Sachiko Arase, and Jun Abe. 2009. “Adaptive Evolution Involving Gene Duplication and Insertion of a Novel Ty1/Copia-like Retrotransposon in Soybean.” Journal of Molecular Evolution 69 (2): 164–75. https://doi.org/10.1007/s00239-009-9262-1.

Kirsten, S., A. Navarro-Quezada, D. Penselin, C. Wenzel, A. Matern, A. Leitner, T. Baum, U. Seiffert, and W. Knogge. 2012. “Necrosis-Inducing Proteins of Rhynchosporium Commune, Effectors in Quantitative Disease Resistance.” Molecular Plant-Microbe Interactions 25 (10): 1314–25. https://doi.org/10.1094/MPMI-03-12-0065-R.

Lamari, Lakhdar. 2002. “Assess: Image Analysis Software for Plant Disease Quantification.” Langmead, Ben, and Steven L. Salzberg. 2012. “Fast Gapped-Read Alignment with Bowtie 2.” Nature Methods 9 (4): 357–59. https://doi.org/10.1038/nmeth.1923.

Langner, Thorsten, and Vera Göhre. 2016. “Fungal Chitinases: Function, Regulation, and Potential Roles in Plant/Pathogen Interactions.” Current Genetics 62 (2): 243–54. https://doi.org/10.1007/s00294-015-0530-x.

Lendenmann, Mark H., Daniel Croll, and Bruce A. McDonald. 2015. “QTL Mapping of Fungicide Sensitivity Reveals Novel Genes and Pleiotropy with Melanization in the Pathogen Zymoseptoria Tritici.” Fungal Genetics and Biology 80: 53–67. https://doi.org/10.1016/j.fgb.2015.05.001.

Leroux, Pierre, and Anne Sophie Walker. 2013. “Activity of Fungicides and Modulators of Membrane Drug Transporters in Field Strains of Botrytis Cinerea Displaying Multidrug Resistance.” European Journal of Plant Pathology 135 (4): 683–93. https://doi.org/10.1007/s10658-012-0105-3.

Lucas, John A., Nichola J. Hawkins, and Bart A. Fraaije. 2015. The Evolution of Fungicide Resistance. Advances in Applied Microbiology. Vol. 90. Elsevier Ltd. https://doi.org/10.1016/bs.aambs.2014.09.001.

Ma, Zhonghua, Tyre J. Proffer, Janette L. Jacobs, and George W. Sundin. 2006. “Overexpression of the 14 α-Demethylase Target Gene (CYP51) Mediates Fungicide Resistance in Blumeriella Jaapii.” Applied and Environmental Microbiology 72 (4): 2581–85. https://doi.org/10.1128/AEM.72.4.2581-2585.2006.

McKenna, Aaron, Hanna Matthew, Eric Banks, Andrey Sivachenko, Kristian Cibulskis, Andrew Kernytsky, Kiran Garimella, et al. 2010. “The Genome Analysis Toolkit: A MapReduce Framework for Analyzing next-Generation DNA Sequencing Data.” Genome Research 20: 1297–1303. https://doi.org/10.1101/gr.107524.110.20.

McLysaght, Aoife, and Daniele Guerzoni. 2015. “New Genes from Non-Coding Sequence: The Role of de Novo Protein-Coding Genes in Eukaryotic Evolutionary Innovation.” Philosophical Transactions of the Royal Society B: Biological Sciences 370 (1678). https://doi.org/10.1098/rstb.2014.0332.

Mérot, Claire, Rebekah A. Oomen, Anna Tigano, and Maren Wellenreuther. 2020. “A Roadmap for Understanding the Evolutionary Significance of Structural Genomic Variation.” Trends in Ecology and Evolution 35 (7): 561–72. https://doi.org/10.1016/j.tree.2020.03.002.

Mishra, Sweta, and Johnathan R. Whetstine. 2016. “Different Facets of Copy Number Changes: Permanent, Transient, and Adaptive.” Molecular and Cellular Biology 36 (7): 1050–63. https://doi.org/10.1128/mcb.00652-15.

Mohd-Assaad, Norfarhan, Bruce A Mcdonald, and Daniel Croll. 2018. “Genome-Wide Detection of Genes Under Positive Selection in Worldwide Populations of the Barley Scald Pathogen.” Genome Biol. Evol. 5: 1315–32. https://doi.org/10.1093/gbe/evy087.

MohdD Assaad, Norfarhan, Bruce A. McDonald, and Daniel Croll. 2016. “Multilocus Resistance Evolution to Azole Fungicides in Fungal Plant Pathogen Populations.” Molecular Ecology 25 (24): 6124–42. https://doi.org/10.1111/mec.13916.

MohdD Assaad, Norfarhan, Bruce A. McDonald, and Daniel Croll. 2019. “The Emergence of the MultiD species NIP1 Effector in Rhynchosporium Was Accompanied by High Rates of Gene Duplications and Losses.” Environmental Microbiology 21 (8): 2677–95. https://doi.org/10.1111/1462-2920.14583.

Morris, J. Jeffrey, Richard E. Lenski, and Erik R. Zinser. 2012. “The Black Queen Hypothesis: Evolution of Dependencies through Adaptive Gene Loss.” MBio 3 (2). https://doi.org/10.1128/mBio.00036-12.

Morschhäuser, Joachim. 2016. “The Development of Fluconazole Resistance in Candida Albicans – an Example of Microevolution of a Fungal Pathogen.” Journal of Microbiology 54 (3): 192–201. https://doi.org/10.1007/s12275-016-5628-4.

Oggenfuss, Ursula, Thomas Badet, Thomas Wicker, Fanny E. Hartmann, Nikhil Kumar Singh, Leen Abraham, Petteri Karisto, et al. 2021. “A Population-Level Invasion by Transposable Elements Triggers Genome Expansion in a Fungal Pathogen.” ELife 10. https://doi.org/10.7554/eLife.69249.

Oki, Nobuhiko, Kentaro Yano, Yutaka Okumoto, Takuji Tsukiyama, Masayoshi Teraishi, and Takatoshi Tanisaka. 2008. “A Genome-Wide View of Miniature Inverted-Repeat Transposable Elements (MITEs) in Rice, Oryza Sativa Ssp. Japonica.” Genes and Genetic Systems 83 (4): 321–29. https://doi.org/10.1266/ggs.83.321.

Olson, Maynard V. 1999. “When Less Is More: Gene Loss as an Engine of Evolutionary Change.” Am. J. Hum. Genet 64: 18–23.

Penselin, Daniel, Martin Münsterkötter, Susanne Kirsten, Marius Felder, Stefan Taudien, Matthias Platzer, Kevin Ashelford, et al. 2016. “Comparative Genomics to Explore Phylogenetic Relationship, Cryptic Sexual Potential and Host Specificity of Rhynchosporium Species on Grasses.” BMC Genomics 17 (953). https://doi.org/10.1186/s12864-016-3299-5.

Petersen, Thomas Nordahl, Søren Brunak, Gunnar Von Heijne, and Henrik Nielsen. 2011. “SignalP 4.0: Discriminating Signal Peptides from Transmembrane Regions.” Nature Methods 8 (10): 785–86. https://doi.org/10.1038/nmeth.1701.

Presti, Libera Lo, Daniel Lanver, Gabriel Schweizer, Shigeyuki Tanaka, Liang Liang, Marie Tollot, Alga Zuccaro, Stefanie Reissmann, and Regine Kahmann. 2015. “Fungal Effectors and Plant Susceptibility.” Annual Review of Plant Biology 66: 513–45. https://doi.org/10.1146/annurev-arplant-043014-114623.

Pritchard, Jonathan K., Matthew Stephens, and Peter Donnelly. 2008. “Inference of Population Structure Using Multilocus Genotype Data.” Genetics 155: 945–59. https://doi.org/10.1007/s10681-008-9788-0.

Quinlan, Aaron R., and Ira M. Hall. 2010. “BEDTools: A Flexible Suite of Utilities for Comparing Genomic Features.” Bioinformatics 26 (6): 841–42. https://doi.org/10.1093/bioinformatics/btq033.

Ritz, Christian, Florent Baty, Jens C. Streibig, and Daniel Gerhard. 2015. “Dose-Response Analysis Using R.” PLoS ONE 10 (12). https://doi.org/10.1371/journal.pone.0146021.

Robinson, Mark D., Davis J. McCarthy, and Gordon K. Smyth. 2009. “EdgeR: A Bioconductor Package for Differential Expression Analysis of Digital Gene Expression Data.” Bioinformatics 26 (1): 139–40. https://doi.org/10.1093/bioinformatics/btp616.

Rohe, Matthias, Angela Gierlich, Hanno Hermann, Matthias Hahn, Bernhard Schmidt, Sabine Rosahl, and Wolfgang Knogge. 1995. “The Race-Specific Elicitor, NIP1, from the Barley Pathogen, Rhynchosporium Secalis, Determines Avirulence on Host Plants of the Rrs1 Resistance Genotype.” EMBO Journal 14 (17): 4168–77. https://doi.org/10.1002/j.1460-2075.1995.tb00090.x.

Sánchez-Torres, Paloma. 2021. “Molecular Mechanisms Underlying Fungicide Resistance in Citrus Postharvest Green Mold.” Journal of Fungi 7 (9). https://doi.org/10.3390/jof7090783.

Sánchez-Vallet, Andrea, Simone Fouché, Isabelle Fudal, Fanny E Hartmann, Jessica L Soyer, Aurélien Tellier, and Daniel Croll. 2018. “The Genome Biology of Effector Gene Evolution in Filamentous Plant Pathogens Andrea.” Annual Review of Phytopathology 56: 12–52. https://doi.org/10.1146/annurev-phyto-080516.

Santiago, Néstor, Cristina Herráiz, J. Ramón Goñi, Xavier Messeguer, and Josep M. Casacuberta. 2002. “Genome-Wide Analysis of the Emigrant Family of MITEs of Arabidopsis Thaliana.” Molecular Biology and Evolution 19 (12): 2285–93.https://doi.org/10.1093/oxfordjournals.molbev.a004052.

Schmidt, Sarah M., Petra M. Houterman, Ines Schreiver, Lisong Ma, Stefan Amyotte, Biju Chellappan, Sjef Boeren, Frank L.W. Takken, and Martijn Rep. 2013. “MITEs in the Promoters of Effector Genes Allow Prediction of Novel Virulence Genes in Fusarium Oxysporum.” BMC Genomics 14 (1): 119. https://doi.org/10.1186/1471-2164-14-119.

Schneider, Caroline A., Wayne S. Rasband, and Kevin W. Eliceiri. 2012. “NIH Image to ImageJ: 25 Years of Image Analysis.” Nat Methods 9 (7): 671–75. https://doi.org/10.1007/978-1-84882-087-6_9.

Schürch, Stéphanie, Celeste C. Linde, Wolfgang Knogge, Lee F. Jackson, and Bruce A. McDonald. 2004. “Molecular Population Genetic Analysis Differentiates Two Virulence Mechanisms of the Fungal Avirulence Gene NIP1.” Molecular Plant-Microbe Interactions 17 (10): 1114–25. https://doi.org/10.1094/MPMI.2004.17.10.1114.

Sionov, Edward, Hyeseung Lee, Yun C. Chang, and Kyung J. Kwon-Chung. 2010. “Cryptococcus Neoformans Overcomes Stress of Azole Drugs by Formation of Disomy in Specific Multiple Chromosomes.” PLoS Pathogens 6 (4). https://doi.org/10.1371/journal.ppat.1000848.

Smit, AFA, R Hubley, and Green P. 2015. “RepeatMasker Open-4.0.” 2015. http://repeatmasker.org.

Sorbo, Giovanni Del, Henk Jan Schoonbeek, and Maarten A. De Waard. 2000. “Fungal Transporters Involved in Efflux of Natural Toxic Compounds and Fungicides.” Fungal Genetics and Biology 30 (1): 1–15. https://doi.org/10.1006/fgbi.2000.1206.

Stanke, Mario, Oliver Keller, Irfan Gunduz, Alec Hayes, Stephan Waack, and Burkhard Morgenstern. 2006. “AUGUSTUS: A b Initio Prediction of Alternative Transcripts.” Nucleic Acids Research 34 (WEB. SERV. ISS.): 435–39. https://doi.org/10.1093/nar/gkl200.

Steenwyk, Jacob L., and Antonis Rokas. 2018. “Copy Number Variation in Fungi and Its Implications for Wine Yeast Genetic Diversity and Adaptation.” Frontiers in Microbiology 9 (288). https://doi.org/10.3389/fmicb.2018.00288.

Stefansson, T. S., Y. Willi, D. Croll, and B. A. McDonald. 2014. “An Assay for Quantitative Virulence in Rhynchosporium Commune Reveals an Association between Effector Genotype and Virulence.” Plant Pathology 63 (2): 405–14. https://doi.org/10.1111/ppa.12111.

Stefansson, Tryggvi S., Bruce A. Mcdonald, and Yvonne Willi. 2013. “Local Adaptation and Evolutionary Potential along a Temperature Gradient in the Fungal Pathogen Rhynchosporium Commune.” Evolutionary Applications 6 (3): 524–34. https://doi.org/10.1111/eva.12039.

Stefansson, Tryggvi S., Bruce A. McDonald, and Yvonne Willi. 2014. “The Influence of Genetic Drift and Selection on Quantitative Traits in a Plant Pathogenic Fungus.” PLoS ONE 9 (11). https://doi.org/10.1371/journal.pone.0112523.

Tam, Vivian, Nikunj Patel, Michelle Turcotte, Yohan Bossé, Guillaume Paré, and David Meyre. 2019. “Benefits and Limitations of Genome-Wide Association Studies.” Nature Reviews Genetics 20 (8): 467–84. https://doi.org/10.1038/s41576-019-0127-1.

Tang, Yun Chi, and Angelika Amon. 2013. “Gene Copy-Number Alterations: A Cost-Benefit Analysis.” Cell 152 (3): 394–405. https://doi.org/10.1016/j.cell.2012.11.043.

Thompson, Julie D., Desmond G. Higgins, and Toby J. Gibson. 1994. “CLUSTAL W: Improving the Sensitivity of Progressive Multiple Sequence Alignment through Sequence Weighting, Position- Specific Gap Penalties and Weight Matrix Choice.” Nucleic Acids Research 22 (22): 4673–80. https://doi.org/10.1093/nar/22.22.4673.

Todd, Robert T., and Anna Selmecki. 2020. “Expandable and Reversible Copy Number Amplification Drives Rapid Adaptation to Antifungal Drugs.” ELife 9: 1–33. https://doi.org/10.7554/eLife.58349.

Trapnell, Cole, Lior Pachter, and Steven L. Salzberg. 2009. “TopHat: Discovering Splice Junctions with RNA-Seq.” Bioinformatics 25 (9): 1105–11. https://doi.org/10.1093/bioinformatics/btp120.

Urban, Martin, Alayne Cuzick, James Seager, Valerie Wood, Kim Rutherford, Shilpa Yagwakote Venkatesh, Nishadi De Silva, et al. 2020. “PHI-Base: The Pathogen-Host Interactions Database.” Nucleic Acids Research 48: 613–20. https://doi.org/10.1093/nar/gkz904.

Vega, Byron, Daniele Liberti, Philip F. Harmon, and Megan M. Dewdney. 2012. “A Rapid Resazurin- Based Microtiter Assay to Evaluate QoI Sensitivity for Alternaria Alternata Isolates and Their Molecular Characterization.” Plant Disease 96 (9): 1262–70. https://doi.org/10.1094/PDIS-12-11-1037-RE.

Wellenreuther, Maren, Claire Mérot, Emma Berdan, and Louis Bernatchez. 2019. “Going beyond SNPs: The Role of Structural Genomic Variants in Adaptive Evolution and Species Diversification.” Molecular Ecology 28 (6): 1203–9. https://doi.org/10.1111/mec.15066.

Wells, Jonathan N., and Cédric Feschotte. 2020. “A Field Guide to Eukaryotic Transposable Elements.” Annual Review of Genetics 54: 539–61. https://doi.org/10.1146/annurev-genet-040620-022145.

White, Shawn E, Ledare F Habera, and R Susan. 1994. “Retrotransposons in the Flanking Regions of Normal Plant Genes: A Role for Copia-like Elements in the Evolution of Gene Structure and Expression.” Proc. Natl. Acad. Sci. USA 91: 11792–96.

Wicker, Thomas, François Sabot, Aurélie Hua-Van, Jeffrey L. Bennetzen, Pierre Capy, Boulos Chalhoub, Andrew Flavell, et al. 2007. “A Unified Classification System for Eukaryotic Transposable Elements.” Nature Genetics 8: 973–82. https://doi.org/10.2118/36071-JPT.

Yike, Iwona. 2011. “Fungal Proteases and Their Pathophysiological Effects.” Mycopathologia 171 (5): 299–323. https://doi.org/10.1007/s11046-010-9386-2.

Yin, Yanbin, Xizeng Mao, Jincai Yang, Xin Chen, Fenglou Mao, and Ying Xu. 2012. “DbCAN: A Web Resource for Automated Carbohydrate-Active Enzyme Annotation.” Nucleic Acids Research 40 (W1): 445–51. https://doi.org/10.1093/nar/gks479.

Zhang, Jingxiang, Liping Li, Quanzhen Lv, Lan Yan, Yan Wang, and Yuanying Jiang. 2019. “The Fungal CYP51s: Their Functions, Structures, Related Drug Resistance, and Inhibitors.” Frontiers in Microbiology 10 (April). https://doi.org/10.3389/fmicb.2019.00691.

