## Supplementary Figures for "The population genetics of adaptation through copy-number variation in a fungal plant pathogen"

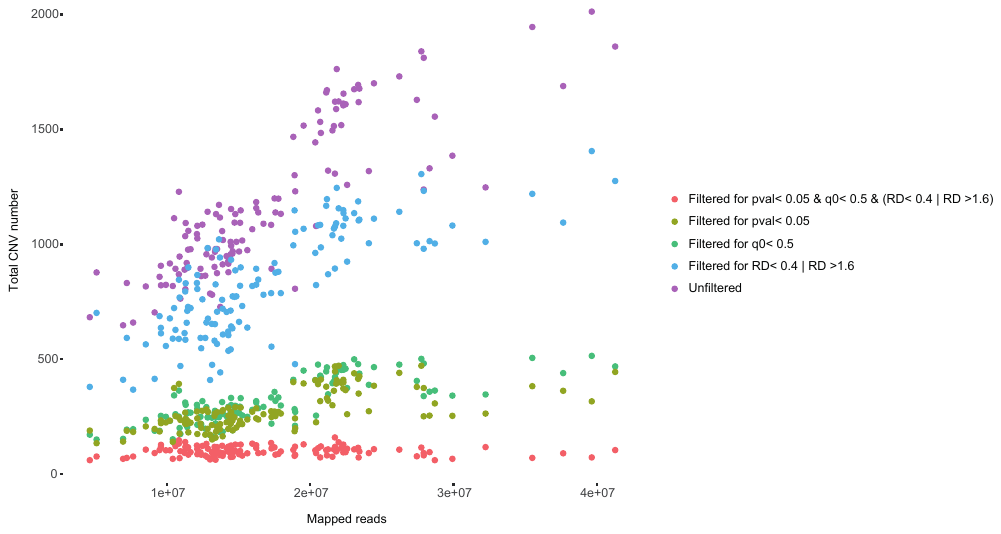

**Supplementary Fig. S1**: For each isolate, total number of CNV calls and the number of CNV calls retained after different filtering steps are shown in relation to total number of mapped reads. After stringent filters on CNV calls, i.e. p-value < 0.05 and q0 < 0.5 and a normalized average read depth of either < 0.4 or > 1.6, there is no correlation of total recovered CNV number and total number of mapped reads.

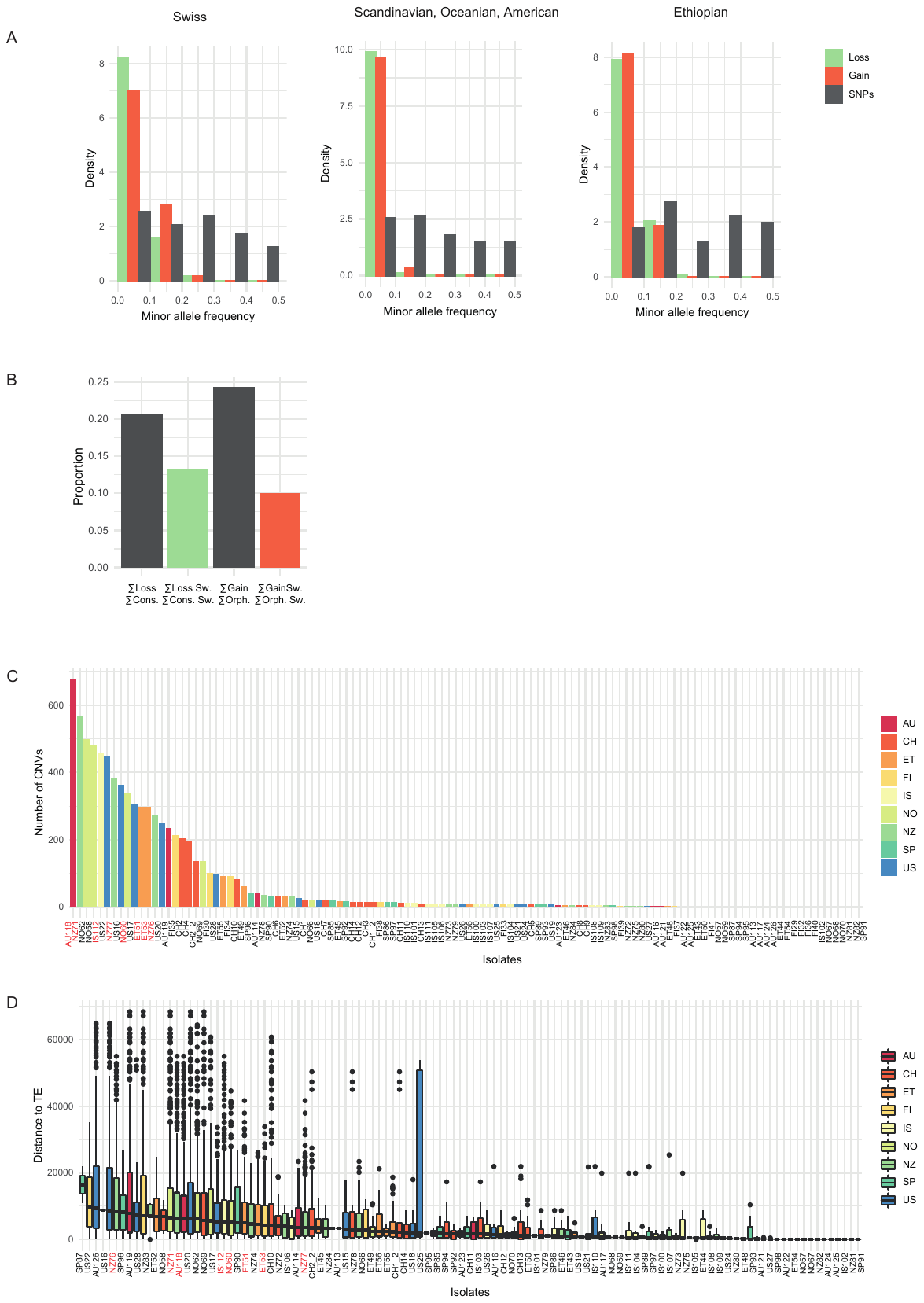

**Supplementary Fig. S2**: (A) Minor allele frequency spectra of loss and gain loci within the three genetic clusters. Spectra were contrasted with the allele frequency of the derived allele at synonymous SNPs. (B) Enrichment analysis of CNV-genes in sweep regions. The number of loss genes divided by the number of conserved genes is compared to the number of sweep genes in loss genes divided by the number of sweep genes in conserved genes. The number of gain genes divided by the number of orphan genes is compared to the number of sweep genes in loss genes divided by the number of sweep genes in orphan genes (Fisher-test loss-conserved p-value < 0.001, gain-orphan p-value < 0.001, n_swe_ in conserved genes = 541, in loss genes = 72, in orphan genes = 9, in gain genes = 90). (C) Total number of CNVs and (D) minimal distance to TEs are shown for each isolate. Colors indicate the country of origin. Isolates in red are differentiated from the main cluster along PC1in the CNV PCA analysis (Fig. 1B).

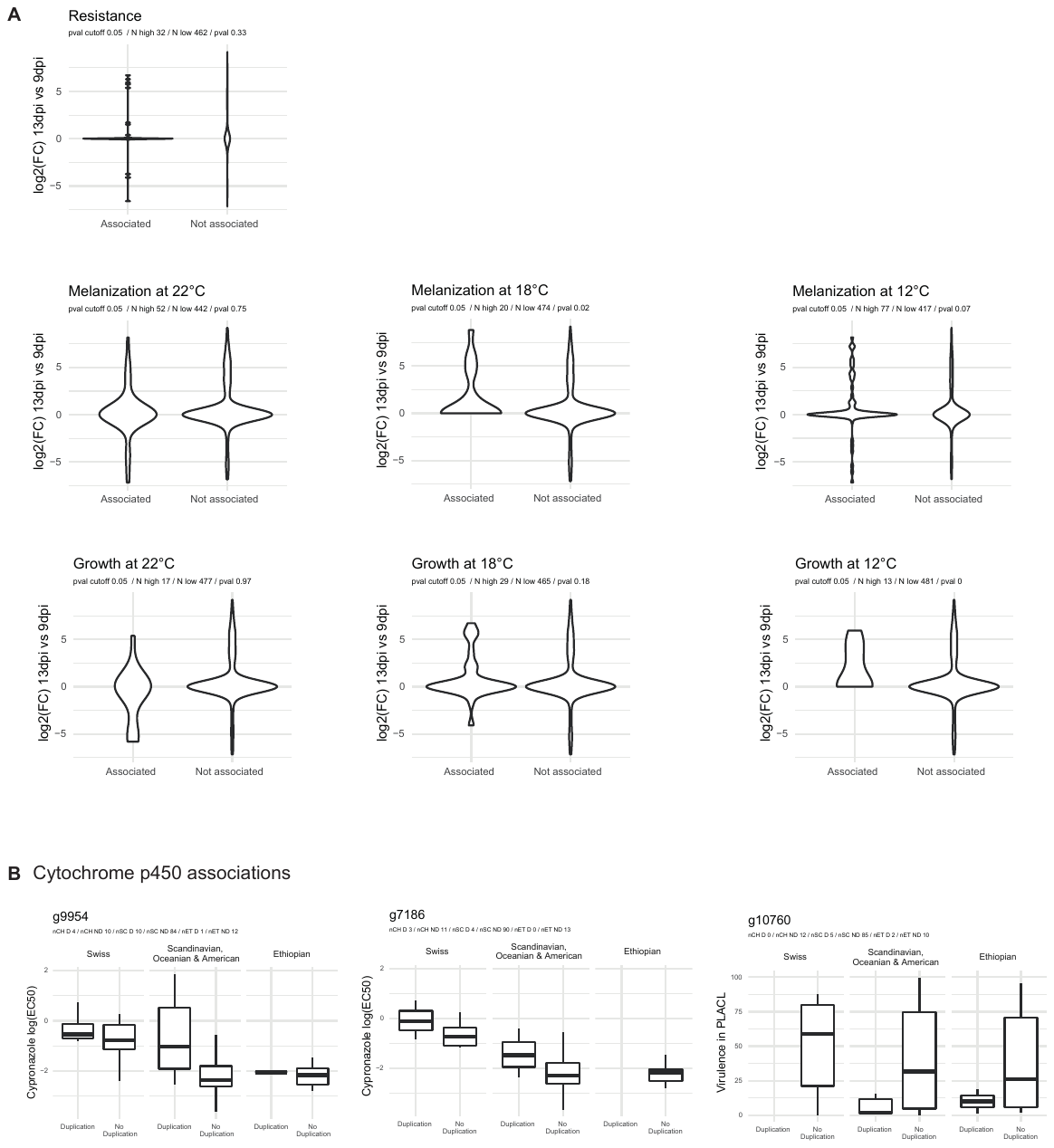

**Supplementary Fig. S3**: (A) Same analysis as in Fig. 2B, but for all the other phenotypes, i.e. cypronazole resistance (measured as EC50), melanization at 12°C, 18°C and 22°C, growth at 12°C, 18°C and 22°C and temperature resilience (defined as standard deviation of the growth rate at 12°C, 18°C and 22°C). (B) Same analysis as Fig. 2D for all significantly associated cytochrome p450 CNV-genes.

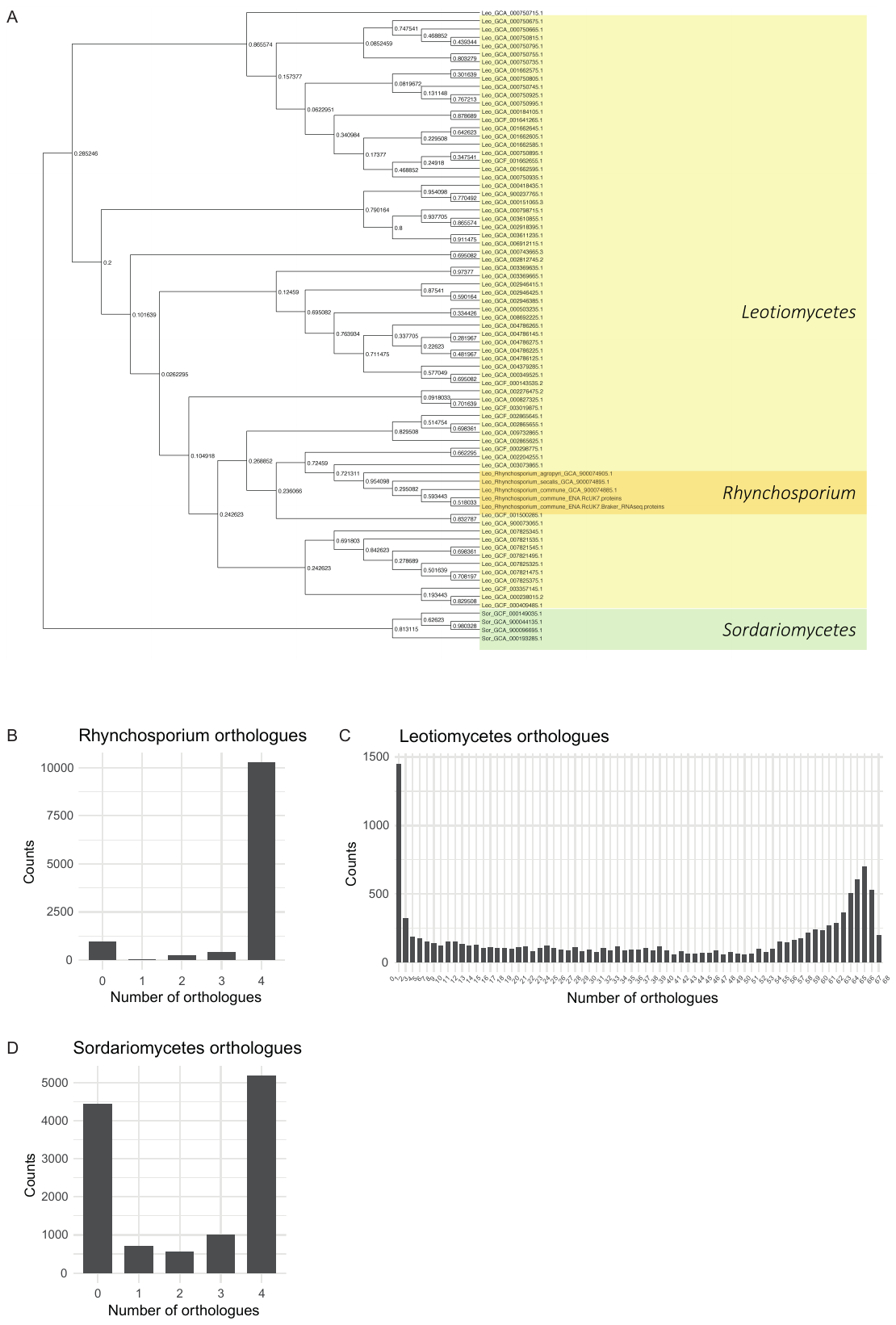

**Supplementary Fig. S4**: (A) Phylogenetic tree of all genomes used for orthologue prediction produced by OrthoFinder. For each *R. commune* gene, the number of orthologs within the analyzed species from the Rhynchosporium genus (A), as well from the Leotimycetes (B) and Sordariomycetes (C) family are shown. The bimodal shape indicates that *R. commune* genes tend to be either conserved within the genus/family or not.

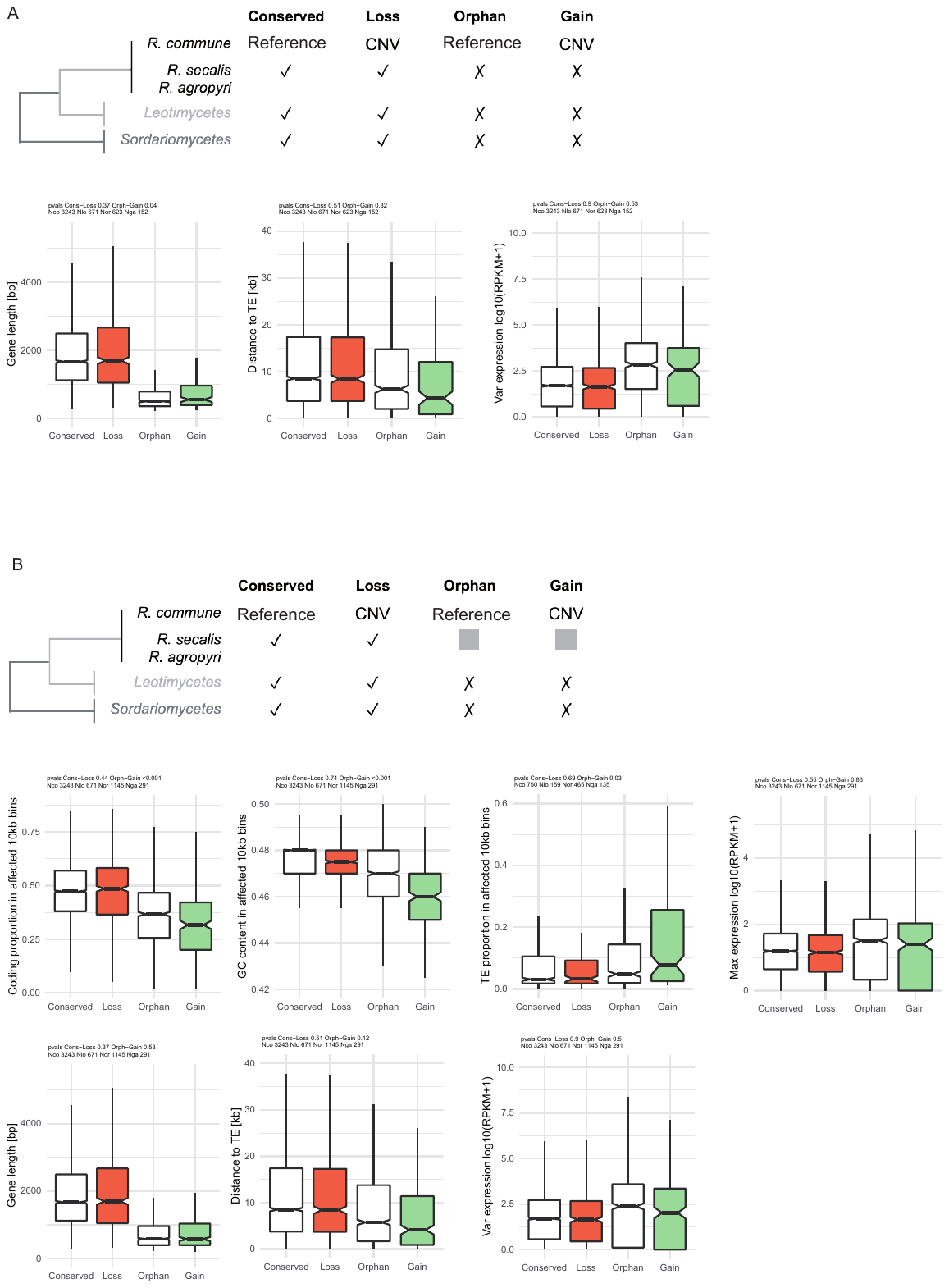

**Supplementary Fig. S5**: (A) Gene length, distance to closest TE and variation in expression is shown for conserved, orphan, loss and gain genes. (B) Same analysis as Fig. 3C-F and Supplementary  Fig.  S5 A, but instead of a species-specific orphan gene definition we used a broader genus-specific definition. Here, orphan genes were defined as genes with no ortholog in Leotimycetes or Sordariomycetes.

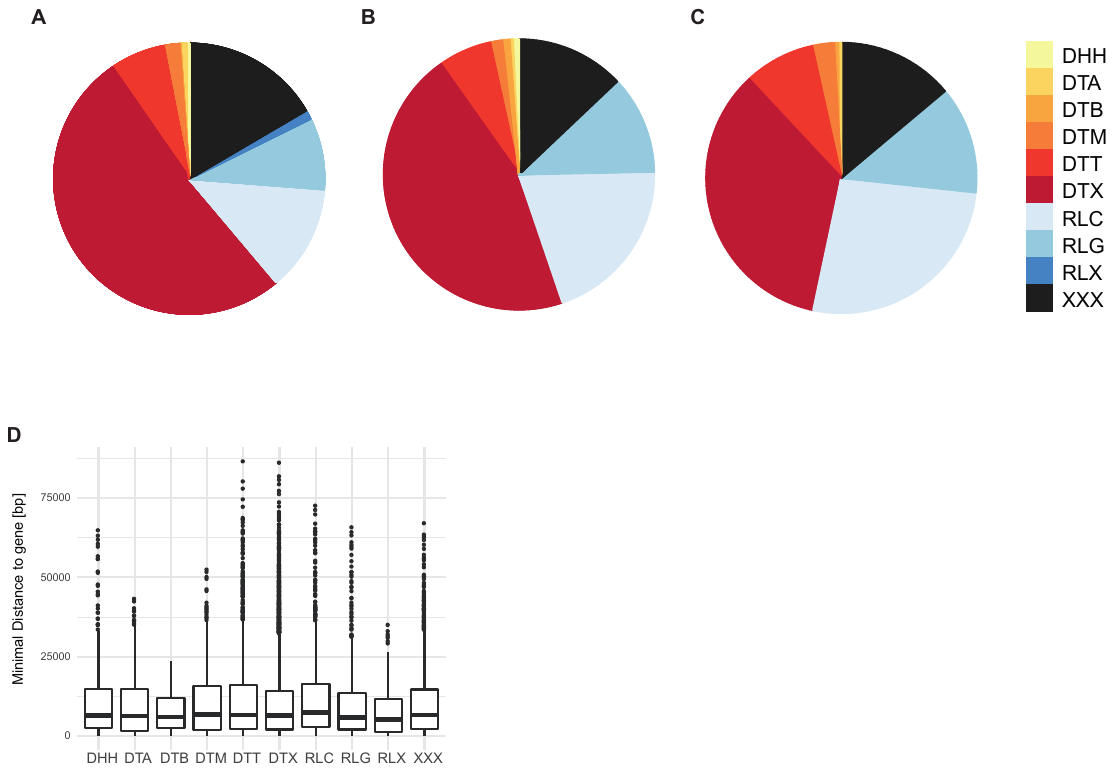

**Supplementary Fig. S6**: (A)-(C) Frequency of TE superfamilies that are closest to genes. Considered is the closest TE for each gene. In (A), all genes were considered, in (B) genes affected by CNVs and in (C) genes affected by a CNV that is significantly associated with at least one of the nine tested phenotypes with an FDR p-value <0.05. TE superfamilies are denoted as DNA-TE TIR hAT (DTA), DNA-TE TIR PiggyBac (DTB), DNA-TE TIR Mariner (DTT), DNA-TE Helitron (DHH), DNA-TE TIR Mutator (DTM), DNA-TE TIR (DTX), Retrotransposon LTR Copia (RLC), Retrotransposon LTR Gypsy (RLG), Retrotransposon LTR (RLX), unknown (XXX). (B) Minimal distance to genes per TE superfamily.

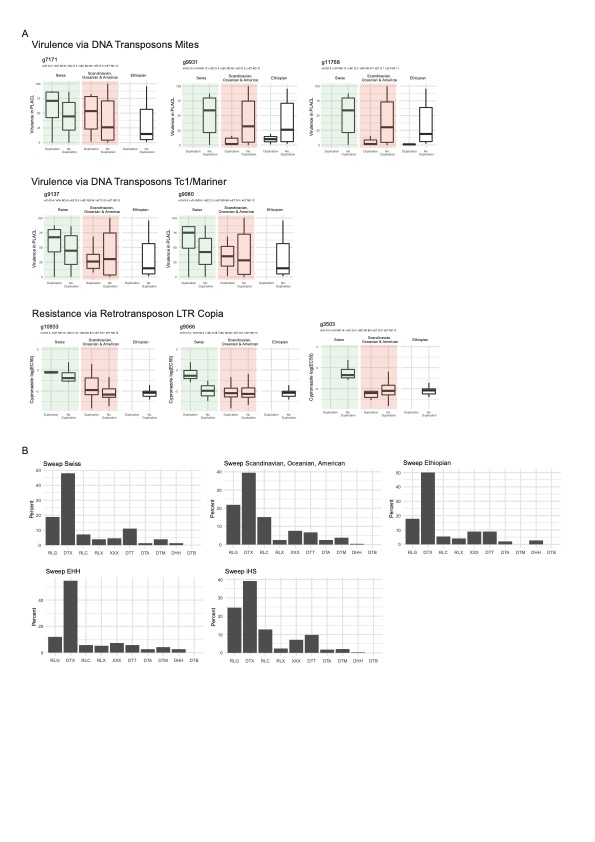

**Supplementary Fig. S7**: (A) Same analysis as Fig. 4C-E. Shown are additional examples of associated CNV-genes close to DTT, DTX or RLC TEs. (B) TE superfamily distribution in sweep regions per genetic cluster, and per statistical determination method.

**Supplementary tables S1-S6**

See separate file “Supplementary_tables.xlsx”
